# The Impact of Neuron Morphology on Cortical Network Architecture

**DOI:** 10.1101/2020.11.13.381087

**Authors:** Daniel Udvary, Philipp Harth, Jakob H. Macke, Hans-Christian Hege, Christiaan P.J. de Kock, Bert Sakmann, Marcel Oberlaender

**Affiliations:** Max Planck Group: In Silico Brain Sciences, Center of Advanced European Studies and Research (caesar), Ludwig Erhard Allee 2, 53175 Bonn, Germany; Department of Visual and Data-centric Computing, Zuse Institute Berlin, Takustraße 7, 14195 Berlin, Germany; Machine Learning in Science, Tübingen University, Maria von Linden Straße 6, 72076 Tübingen, Germany; Department of Integrative Neurophysiology, Center for Neurogenomics and Cognitive Research, VU Amsterdam, De Boelelaan 1085, 1081 Amsterdam, the Netherlands; Max Planck Institute of Neurobiology, Am Klopferspitz 18, 82152 Martinsried, Germany

**Author notes:** **Editorial correspondence:** Max Planck Group: In Silico Brain Sciences, Center of Advanced European Studies and Research (caesar), Ludwig-Erhard-Allee 2, Bonn, 53175 Germany.

## Abstract

It has become increasingly clear that the neurons in the cerebral cortex are not randomly interconnected. This wiring specificity can result from synapse formation mechanisms that interconnect neurons depending on their activity or genetically defined identity. Here we report that in addition to these synapse formation mechanisms, the structural composition of the neuropil provides a third prominent source by which wiring specificity emerges in cortical networks. This structurally determined wiring specificity reflects the packing density, morphological diversity and similarity of the dendritic and axonal processes. The higher these three factors are, the more recurrent the topology of the networks. Conversely, low density, diversity and similarity yield feedforward networks. These principles predict connectivity patterns from subcellular to network scales that are remarkably consistent with empirical observations from a rich body of literature. Thus, cortical network architectures reflect the specific morphological properties of their constituents to much larger degrees than previously thought.

## Introduction

Neuronal networks are implemented in the brain via a plethora of molecular mechanisms, which form synaptic connections between the neurons’ long and branched dendrites and axons (*1-5*). Axo-dendritic proximity is hence necessary for the formation of synapses and constrains which neurons could in principle be connected – and where along their neuronal processes these connections could occur. Yet it is unclear, to what degree the highly diverse morphological properties of the neurons can impact the architecture of the networks they form (*6*). For the past decades, the strategy for investigating the impact of neuron morphology on network architecture has been to test the validity of ‘Peters’ rule’ (*7*). According to this longstanding hypothesis, axons form synaptic connections randomly wherever they get in close proximity to a dendrite (*8*). Proximity would hence not only be necessary, but sufficient to account for connectivity. Consequently, Peters’ rule can be simply restated as “ proximity predicts connectivity”. If this were true, the morphological properties of neurons would directly determine the architecture of neuronal networks, independent of the particular mechanisms that form synapses.

Tests of Peters’ rule have failed to support it (*9-13*). At subcellular scales, dense reconstructions in mouse somatosensory cortex showed that on average, dendritic spines are in close apposition to ten axonal branches, of which generally only one formed a synapse (*12*). Moreover, a subset of these axonal branches formed clusters of synapses by connecting to multiple close-by spines along the same dendrite (see also (*13*)). At cellular scales, sparse reconstructions showed that axo-dendritic proximity is insufficient to account for the patterns of synaptic connections (*14*), or for differences in connection probabilities that reflect the neurons’ cell types (*11*). Because they are inconsistent with Peters’ rule, such observations are considered to reflect wiring principles that are independent of the morphological properties of neurons. Thus, the absence of synaptic connections between close-by dendrites and axons, and the occurrence of clusters of synapses between them, are interpreted as the result of synapse formation mechanisms that connected these neurons based on their cellular identity or activity (*12*).

Interpreted as further evidence for this conclusion are observations which showed that neurons form particular network motifs – for example feedforward loops – that occur more (or less) frequently than expected for randomly connected networks (*15, 16*). Theoretically, such nonrandom topological properties could very well arise from simple sets of wiring rules (*4, 5*) or via learning (*17*), and therefore be independent of the morphological properties of neurons. However, even if neurons were interconnected randomly by axo-dendritic proximity, the resulting networks would also display complex connectivity patterns (*18, 19*) and generally nonrandom topologies (*19, 20*). Furthermore, morphological properties – such as dendrite polarity – can constitute a defining source for nonrandom occurrences of network motifs (*21*). These recent studies indicate that inconsistency with Peters’ rule may be insufficient to conclude whether empirically observed wiring specificity reflects the neurons’ morphology, a particular synapse formation mechanism, or a combination thereof. Thus, conclusive answers to the questions – Which principles link neuron morphology to network architecture, and how can these principles be disentangled from genetically encoded and activity-dependent wiring specificity? – remain presently unknown.

Here, we quantitatively address these questions by utilizing a structural model of the rat barrel cortex (reviewed in (*22*)). The construction of the structural model, including a comprehensive description of the underlying assumptions and anatomical data, was reported previously (*19*) and is briefly summarized in the **Materials and Methods**. Here, we combine the structural model with a statistical approach to derive all configurations of how networks could appear in barrel cortex if synapse formation mechanisms that introduce wiring specificity were absent. In comparison to Peters’ rule, our approach considers axo-dendritic proximity solely as a necessary condition for synapse formation. We demonstrate that proximity can predict connectivity for only a very small minority of the neurons. The impact of neuron morphology on network architecture is hence not direct such that it would determine which neurons are interconnected. Instead, we find that neuron morphology impacts network architecture indirectly by shaping the structural composition of the neuropil – and it is the specific neuropil structure that translates into specific pairwise and higher-order connectivity patterns. We show that the neuropil structure predicts cell type-specific connectivity patterns, occurrences of clusters of synapses, as well as of nonrandom network motifs that are in remarkable agreement with the available empirical data. Finally, we apply our statistical approach to a dense reconstruction of a petascale volume of human cortex (*23*) to illustrate how the impact of neuron morphology on network architecture can be tested beyond the structural model that is used to derive it.

## Results

We previously introduced a reverse engineering approach to construct an anatomically detailed model, which mimics the structural composition of the neuropil for the volume of rat barrel cortex that represents the 24 major facial whiskers (*19*). This structural model is based on quantitative anatomical data that we had systematically collected for the rat barrel cortex and primary thalamus of the whisker system (i.e., the ventral posterior medial nucleus (VPM)), and which we briefly review in the **Materials and Methods**. Here we show that the structural model provides realistic and robust estimates for the amounts of axonal and dendritic branches within any subvolume of the barrel cortex, for the numbers of presynaptic (i.e., boutons) and postsynaptic structures (e.g. spines) that these branches represent, and for the locations and cell types of the neurons from which these branches originate (**Fig. S1** and **Supplementary Text**).

Although the structural model lacks connectivity, it allows us to explore how the morphological properties of the neurons could in principle impact network architectures in the barrel cortex (*19*). For this purpose, we divide the structural model into subvolumes and generate different network configurations that account for all pre- and postsynaptic structures therein **(Fig. S2)**. We assume that boutons of excitatory neurons can connect to the spines from all other excitatory neurons, as well as to postsynaptic structures on the somata and dendritic shafts of all inhibitory neurons (*19*). Boutons of inhibitory neurons can connect to postsynaptic structures on the somata and dendritic shafts of all excitatory and inhibitory neurons, but not to spines. We illustrate our strategy for a schematic subvolume that comprises three boutons along one axonal branch and nine spines along the dendritic branches from four excitatory neurons **(Fig. 1A)**. When each of the boutons is assigned to one of the spines, the structural composition of this particular subvolume alone gives rise to 9!/(9 − 3)! = 504 network configurations that differ in how the five neurons from which these branches originate could be connected to one another **(Fig. S2A)**.

**Figure 1.**
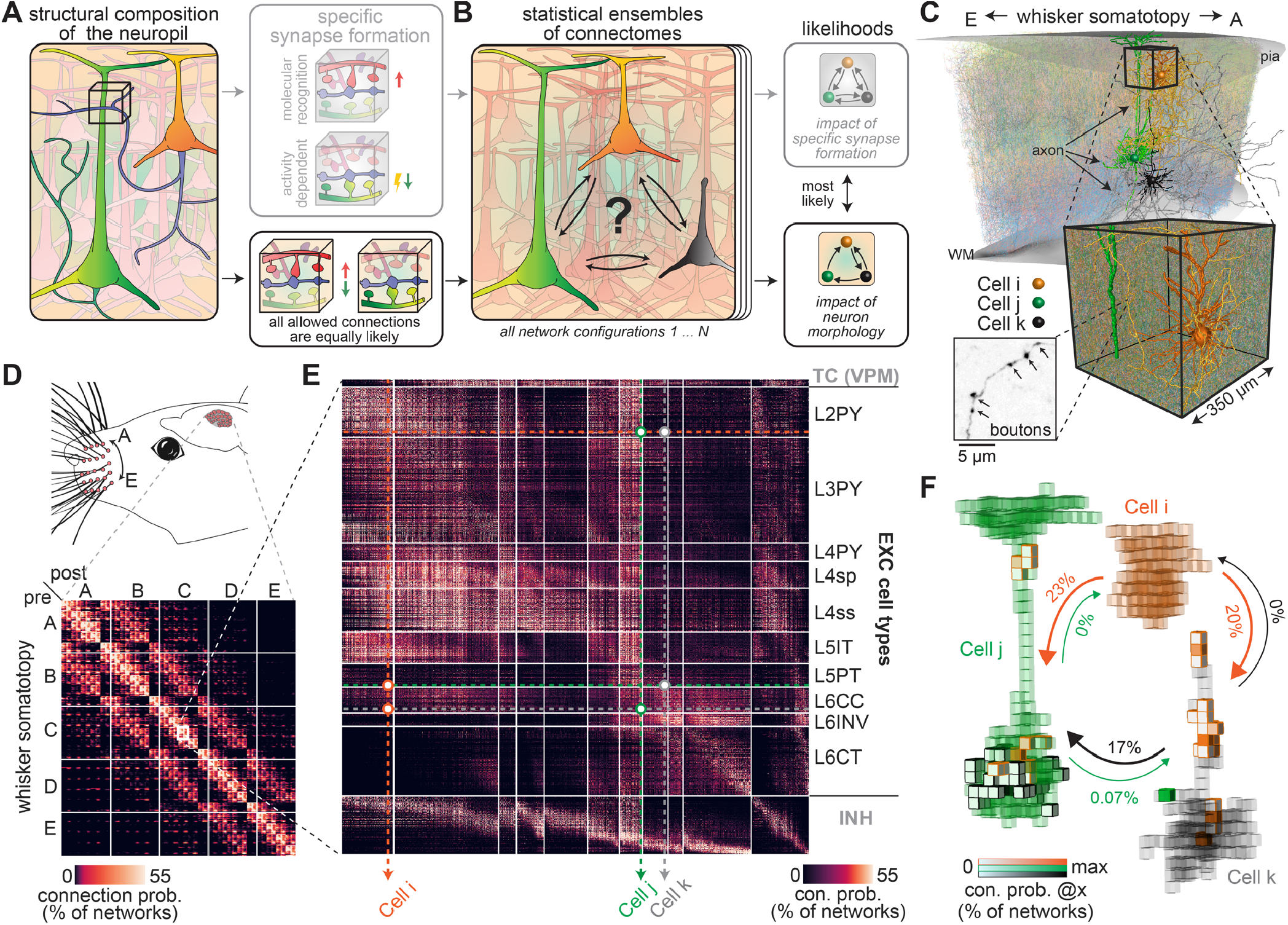
Concept for testing the impact of neuron morphology on network architecture. **(A)** Schematic illustration: We divide the neuropil into small subvolumes, count the presynaptic (i.e., boutons) and postsynaptic structures (e.g. spines) along axons and dendrites, and generate different networks which can account for them. Here, each of the three axonal boutons (blue) could be connected to one of the nine spines from four different neurons (green, red, pink, purple), resulting in 504 different network configurations **(Fig. S2A). (B)** The resulting statistical ensemble of connectomes allows us to calculate how likely each of such network configurations would appear in the absence of synapse formation mechanisms that result in wiring specificity (e.g. molecular recognition and activity dependence). **(C)** Illustration of the structural model of the rat barrel cortex **(Fig. S1)**. The structural model is based on axon and dendrite morphologies of *in vivo* labeled neurons, including distributions of pre- (i.e., boutons) and postsynaptic structures (e.g. spines) along these neuronal processes. Three example neurons from layers 2, 5 and 6 are highlighted. Inset illustrating how boutons were quantified is adapted from (*19*). **(D)** Matrix representation of the statistical ensemble of connectomes for the cortical volume that represents the 24 major facial whiskers (i.e., A1-E4, α-δ). Connection probabilities reflect the fraction of networks in which the respective neurons are connected. **(E)** Zoom-in to the barrel column representing the C2 whisker. The matrix is sorted by soma depth and cell type: VPM (only presynaptic), INH and EXC (subdivided into pyramidal neurons in layers 2-4 (L2PY, L3PY, L4PY), star pyramids (L4sp), spiny stellates (L4ss), intratelencephalic (L5IT), pyramidal tract (L5PT), corticocortical (L6CC), corticothalamic (L6CT) and inverted (L6INV) neurons). **(F)** Axo-dendritic overlap between the neurons from panels C/E (here at 50 µm resolution). Arrows denote the likelihoods that e.g. the axon of *Cell i* is connected by at least one synapse to the dendrites of *Cell j* within one of their overlap volumes (i.e., in 23% of the networks from the statistical ensemble of connectomes).

We calculate how likely each network configuration would occur, when synapse formation mechanisms that result in wiring specificity are neglected (e.g. molecular recognition and activity dependence). For this purpose, we assume that all boutons are equally likely to connect independently to any of the allowed postsynaptic structures within the same subvolume **(Fig. S2B/C)**. These wiring rules (Equations 1-3 in the **Materials and Methods**) hence provide the likelihoods for how neuronal networks could appear if they were solely due to the underlying structural composition of the neuropil **(Fig. 1B)**. Analogous to statistical mechanics (*24*), we refer to the resulting probability distribution of network configurations as a ‘statistical ensemble of connectomes’. Using this strategy, we derive all network configurations that can emerge from the structural model of the barrel cortex **(Fig. 1C)**, calculate how likely these networks are to appear in the absence of synapse formation mechanisms that introduce wiring specificity **(Fig. 1D)**, and analyze the resulting statistical ensemble of connectomes with respect to the neurons’ cell types and locations **(Fig. 1E)**. Thus, we provide quantitative predictions for how the structural composition of the neuropil could impact wiring between arbitrarily grouped neurons across the barrel cortex, and the likelihoods for where along their dendrites and axons connections could occur **(Fig. 1F)**.

### Peters’ rule could predict connectivity for <1% of the neurons

We use the statistical ensemble of connectomes to test whether the absence of synapse formation mechanisms that introduce wiring specificity would result in networks that obey Peters’ rule. For this purpose, we determine the axo-dendritic overlap at different spatial resolutions (here: cubes with 1, 5, 10, 25, 50 or 100 µm edge lengths) for all neuron pairs and quantify the respective amounts of pre- and postsynaptic structures that these neuronal processes represent within each overlap volume **(Fig. 2A)**. We illustrate these quantifications for one example subvolume, which contains more than 100,000 boutons and postsynaptic structures, respectively **(Fig. 2B)**. When each of the boutons is assigned to one of the allowed postsynaptic structures, the composition of this particular subvolume alone gives rise to an enormous number of network configurations that differ in how the ∼15,000 neurons from which these synaptic structures originate are interconnected **(Fig. S2B)**. In the vast majority of these networks, any particular neuron pair remains unconnected – i.e., their pre- and postsynaptic structures connect to those of other neurons that overlap in this subvolume **(Fig. 2C)**. Moreover, most neurons contribute more than one synaptic structure to this subvolume, which results in configurations where axon-dendrite pairs are interconnected by several synapses. In any network configuration, and across all subvolumes, 99.6 ± 0.1% of the axon-dendrite pairs remain unconnected despite their overlap, while a substantial fraction of them forms clusters with up to five (or more) synapses **(Fig. 2D)**. Thus, axo-dendritic overlap could predict connectivity, but in principle for only a very small minority of overlapping branch pairs. This general deviation from Peters’ rule is due to the high packing density of neuronal processes, which leads to a number of axon-dendrite pairs that exceeds the number of synapses that they represent by one to two orders of magnitude, irrespective of the spatial resolution at which overlap is determined **(Fig. 2E)**.

**Figure 2.**
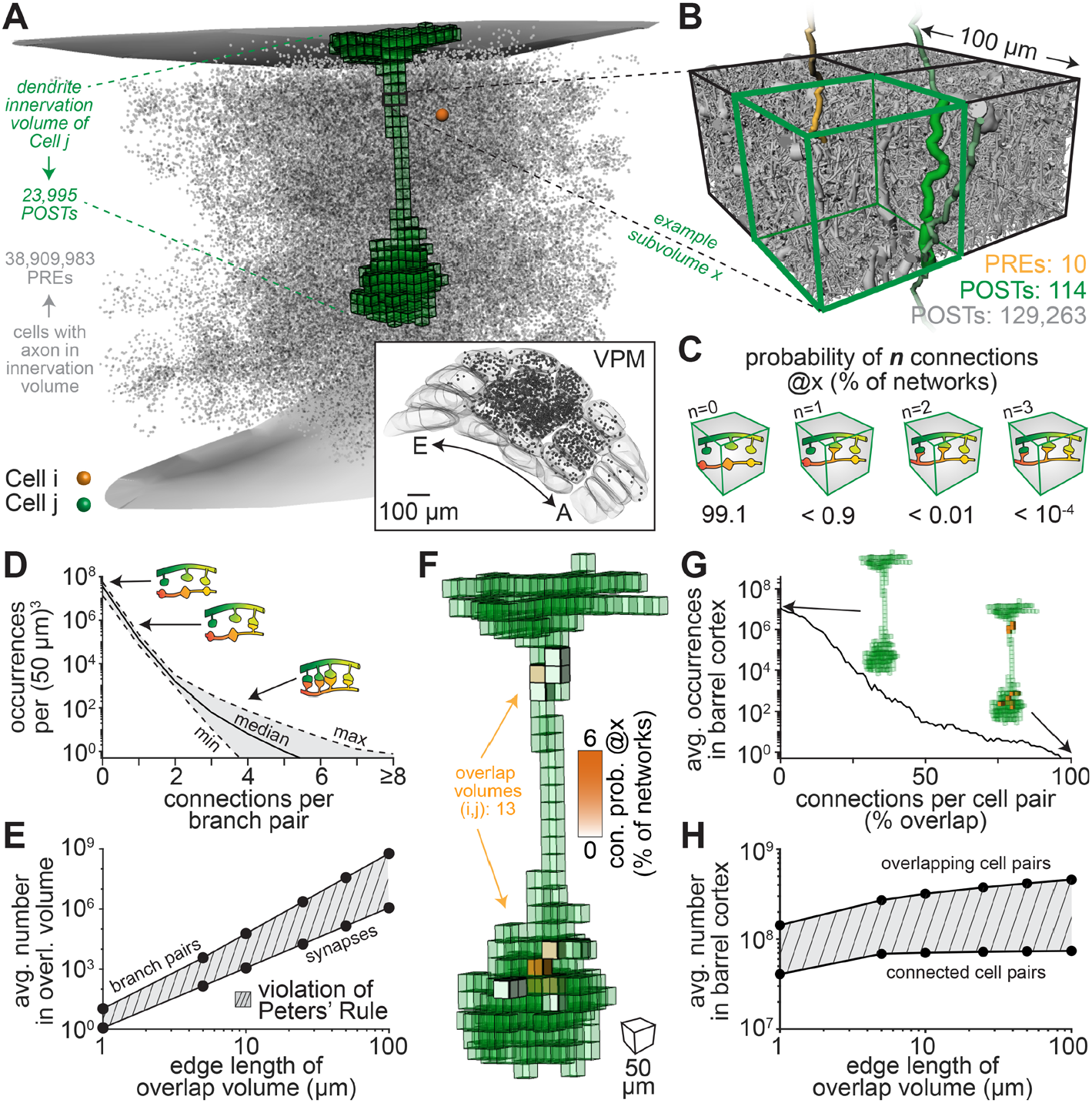
Testing of Peters’ rule. **(A)** Neurons (grey, 50% shown) in barrel cortex and VPM thalamus whose axons overlap (here at 50 µm resolution) with the L5PT dendrites from **Fig. 1F. (B)** Subvolume in which the L2PY axon from **Fig. 1F** represents ten presynaptic structures (PREs; i.e., boutons). The L5PT dendrites represent 114 postsynaptic structures (POSTs; i.e., spines). The number of 129,263 POSTs reflects all spines in this subvolume, and postsynaptic structures on inhibitory somata and dendrites **(Fig. S2B). (C)** Probabilities that none, one or more of the pre- and postsynaptic structures of the L2PY and L5PT are connected in the statistical ensemble of connectomes **(Fig. S2C). (D)** Axon-dendrite pairs whose overlap yields none, one or more connections across all networks from the statistical ensemble of connectomes. **(E)** Axon-dendrite pairs versus the number synapses that they represent for different resolutions at which axo-dendritic overlap is determined. If the statistical ensemble of connectomes were consistent with Peters’ rule, the two lines would match. **(F)** The L2PY axon overlaps with the L5PT dendrites in 13 subvolumes. **(G)** Number of overlap volumes for all neuron pairs versus connections between them. 100% indicates that two neurons are connected by as many synapses as they have overlap volumes. **(H)** Neuron pairs that overlap versus connected neurons across networks from six statistical ensembles of connectomes that differ in the resolution at which overlap is determined.

The deviations from Peters’ rule become even larger at cellular scales. The axons and dendrites of neuron pairs typically overlap in more than one subvolume **(Fig. 2F)**. The probability that overlap could predict connectivity is already very low within each individual subvolume. The likelihood that networks occur in which any particular neuron pair is interconnected in all of its overlap volumes is hence in general infinitesimally small **(Fig. 2G)**. Irrespective of the spatial resolution at which overlap is determined, a vast majority of the neurons whose axons and dendrites overlap will remain unconnected in any of the networks that could emerge from the structural composition of the barrel cortex’ neuropil **(Fig. 2H)**. Thus, empirical observations that falsify Peters’ rule do in general not provide insight into the underlying synapse formation mechanisms, and in particular do not support conclusions that these neurons must have been interconnected depending on their cellular identity or activity. Consequently, testing the validity of Peters’ rule is insufficient for testing the impact of neuron morphology on network architecture.

### Nonrandom networks can emerge due to the structural composition of the neuropil

Next, we test the degrees to which the absence of synapse formation mechanisms that introduce wiring specify would result in random networks. For this purpose, we compare the topology of networks from the statistical ensemble of connectomes with those of randomly connected networks. Specifically, we calculate the respective occurrences of all fifteen wiring patterns by which three neurons could be connected to one another – commonly referred to as triplet motifs **(Fig. 3A)**. The likelihood that three neurons in the structural model of the barrel cortex form a particular motif depends on their respective pairwise connection probabilities in the statistical ensemble of connectomes. For example, the three neurons shown in **Figure 1F** will most likely form the motif with a single unidirectional connection (i.e., L2PY → L5PT), less likely motifs with one bidirectional connection (i.e., L5PT ←→ L6CC), and never motifs with more than one bidirectional connection (i.e., the L5PT and L6CC axons do not overlap with the L2PY dendrites). Consequently, the motifs that are formed by any particular subset of neurons will differ across the different networks from the statistical ensemble of connectomes **(Fig. 3B)**. However, in any of these networks, the respective occurrences of motifs deviate from those in random networks **(Fig. 3C)**, although both the networks from the statistical ensemble of connectomes and the random networks are based on the same pairwise connection probability distributions **(Fig. 3D)**.

**Figure 3.**
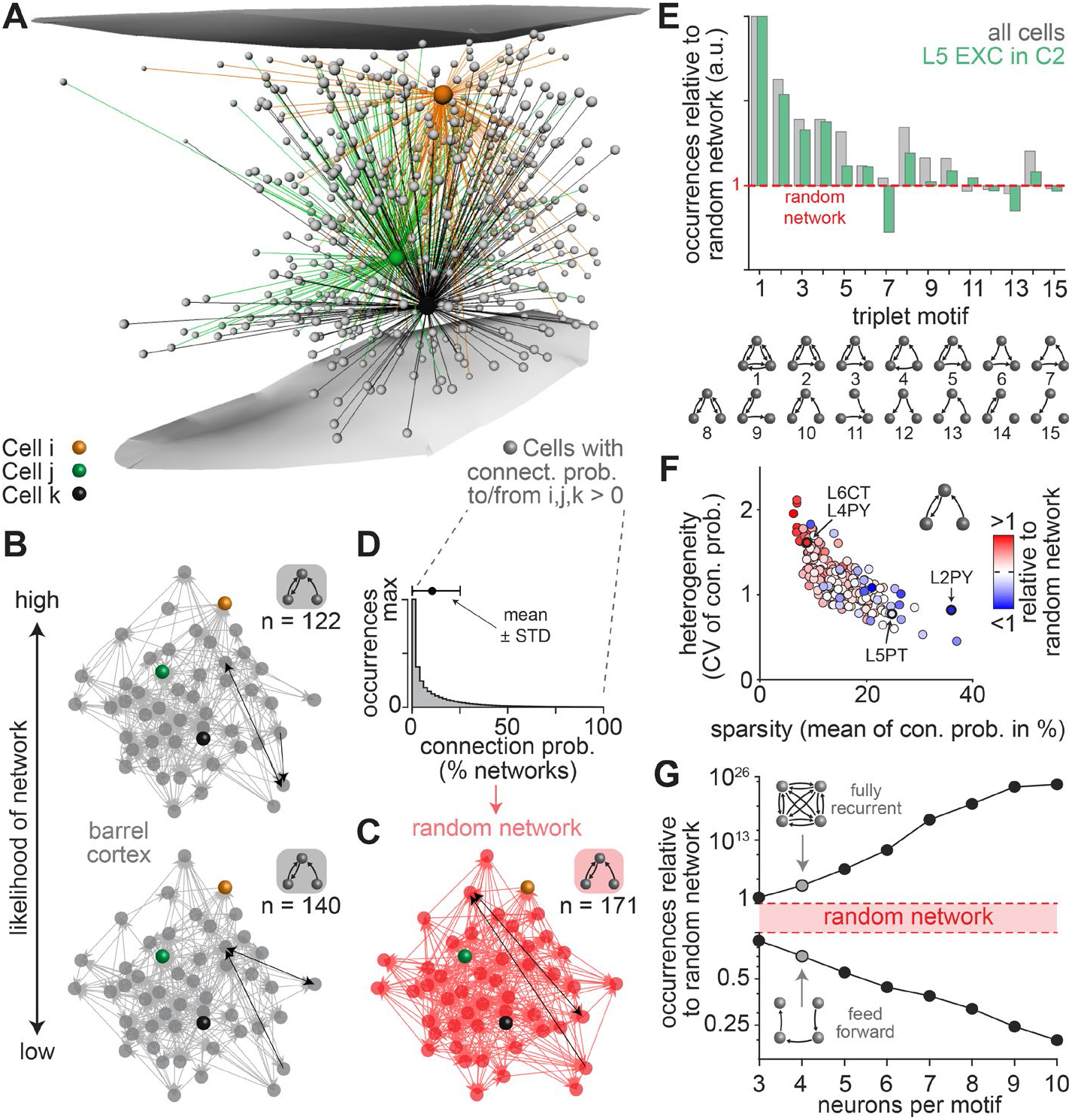
Relationships between neuron morphology and nonrandom network topology. **(A)** Somata of example neurons (grey) whose axons overlap with the dendrites of at least one of the neurons from **Fig. 1F. (B)** Two network configurations from the statistical ensemble of connectomes for 50 neurons from panel A that are predicted to occur with different likelihoods. **(C)** Random network example derived from the same pairwise connection probability distribution that was used to generate the networks in panel B. The number of nodes and edges are identical to panel B, but the occurrences (n) of motifs differ (e.g. motif 10 in panel E). One example for motif 10 is highlighted in black per network in panels B/C. **(D)** Connection probability distribution for all pairs of neurons from panel A. **(E)** Ratios between motif occurrences in networks from the statistical ensemble of connectomes versus random networks (1: equally abundant; >1: overrepresented compared to random networks; <1: underrepresented). **(F)** Deviations in the occurrences of motif 10 compared to random networks for different cell type-specific groupings (n=220) versus the respective means and CVs of the underlying connection probability distributions. **(G)** The likelihoods to observe recurrent loops and feedforward chains between up to ten neurons in networks from the statistical ensemble of connectomes versus in random networks.

The degrees to which the occurrences of motifs differ from those in random networks depends on how the neurons are grouped. For example, while feedforward loops (motif 7) occur generally more frequently in the networks from the statistical ensemble of connectomes than in random networks (i.e., they are overrepresented), they are underrepresented in the subnetworks formed by excitatory neurons in layer 5 **(Fig. 3E)**. The respective occurrences of any of the triplet motifs change depending on how neurons are grouped with respect to layer, barrel column, cell type, inter-somatic distance, or combinations thereof **(Fig. S3)**. Interestingly, these grouping-specific occurrences of motifs correlate with differences in the shapes of the underlying pairwise statistics. For example, the occurrences of the recurrent feedforward motif (motif 10) transition from over- to underrepresentation depending on the respective means of the connection probability distributions **(Fig. 3F)**. The magnitude of over- and underrepresentation increases with increasing and decreasing widths of the connection probability distributions, respectively. These relationships indicate that the mean and the coefficient of variation (CV) of connection probabilities – here referred to as the degrees of network sparsity and heterogeneity – can quantitatively and qualitatively impact the nonrandom topology of neuronal networks **(Fig. 3F)**. Beyond triplet motifs, recurrence characterizes the networks from the statistical ensemble of connectomes, as overrepresentation is predicted to increase with the number of bidirectional connections and with the number of neurons per motif **(Fig. 3G)**. Conversely, underrepresentation of feedforward motifs increases with the number of neurons per motif. Consequently, the topology that emerges in networks due to the structural composition of the neuropil will in general deviate from those of random networks, even if synapse formation mechanisms that introduce wiring specificity were absent. Thus, empirically observed nonrandom motif occurrences do in general not provide insight into the underlying synapse formation mechanisms, and do not support conclusions that such patterns reflect a particular wiring rule.

### Three morphology-related factors translate into nonrandom network architectures

How is it possible that the structural composition of the neuropil translates into nonrandom network architectures? How can the degrees of sparsity and heterogeneity in connectivity have such a defining impact on the networks’ specific nonrandom topological properties? To address these questions, we systematically explore the mathematical basis that underlies the occurrences of network motifs. For this purpose, we consider the statistical ensemble of connectomes as a distribution of pairwise connection probabilities that generates network configurations. If each of the connections is drawn independently from any such generating distribution, motifs will occur as expected for randomly connected networks – i.e., occurrences are independent from the network’s heterogeneity and only reflect the mean of the underlying pairwise statistics **(Fig. 4A**; Equation S1 in the **Supplementary Text)**. Thus, our observations of nonrandom occurrences of motifs, and their dependencies on network heterogeneity, cannot be consistent with the assumption that connection probabilities are independent of one another. Instead, only correlations in the statistical ensemble of connectomes could explain our observations **(Fig. 4B)**.

**Figure 4.**
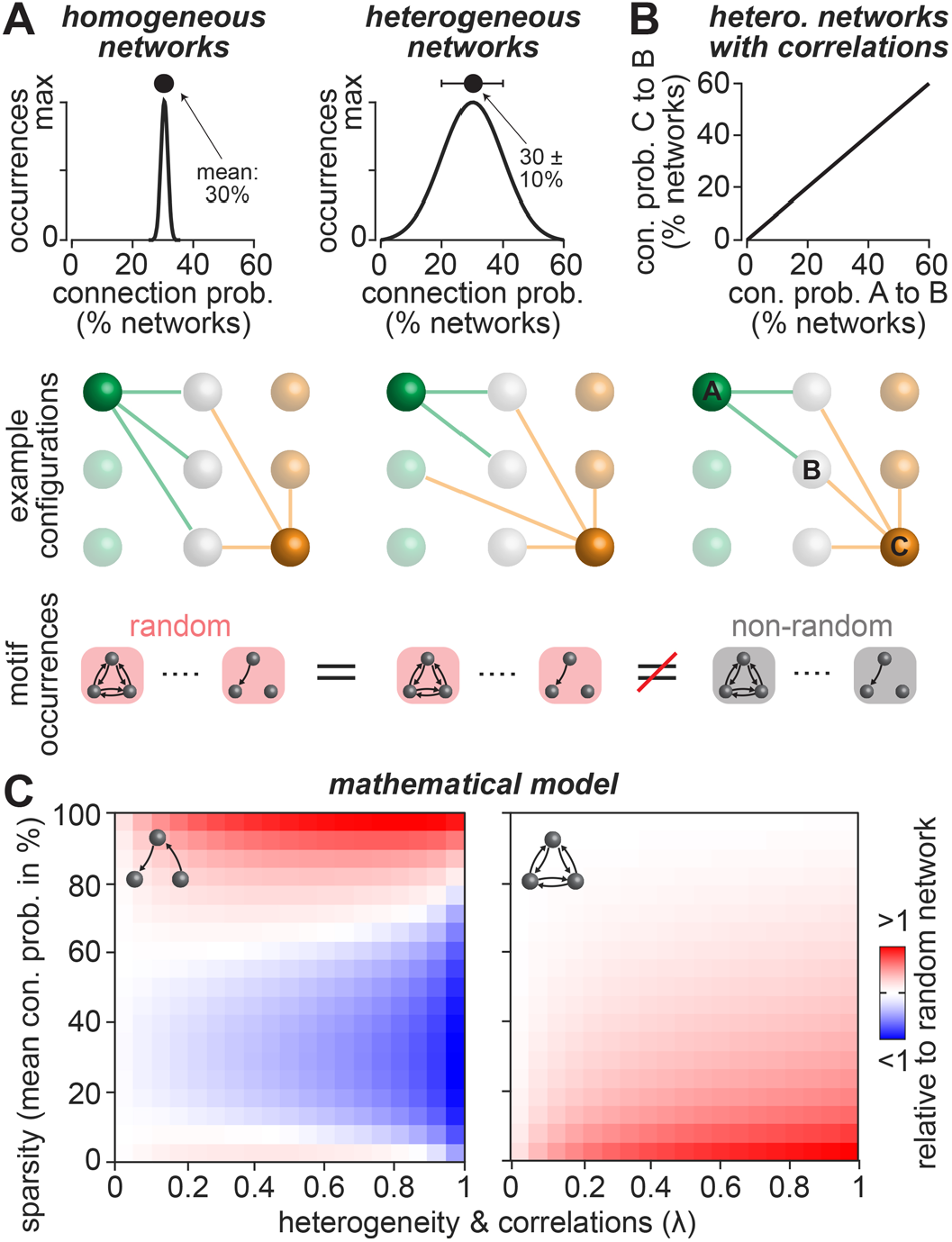
Mathematical basis for nonrandom motif occurrences. **(A)** Toy example illustrating Equation S1 in the **Supplementary Text**. Left: Statistical ensembles of connectomes that yield homogeneous connectivity (i.e., standard deviation << mean) lead to networks in which all nodes have mostly the same number of edges (here illustrated by two nodes that both have three edges). Right: statistical ensembles of connectomes that yield heterogeneous connectivity (i.e., standard deviation ≈ mean) lead to networks in which the nodes have on average the same number of edges. Bottom panels: occurrences of triplet motifs will be on average random and identical in all networks that have the same mean connection probability. **(B)** Toy example illustrating Equations S2-3 in the **Supplementary Text**. Statistical ensembles of connectomes that yield correlated connectivity (here illustrated by two nodes that both connect to the same nodes) will result in motif occurrences that deviate from those of random networks. **(C)** A mathematical model (see **Supplementary Text**) that represents statistical ensembles of connectomes with correlations reveals how the mean (sparsity) and CV (heterogeneity) of pairwise connectivity impact nonrandom motif occurrences; here illustrated for feedforward chains (left) and recurrent loops (right).

To investigate how correlations affect the occurrences of motifs, we developed a mathematical model, which assumes that correlations and heterogeneity in connectivity can be expressed by a single parameter *λ* (*25*). A comprehensive description of the mathematical model is provided in the **Supplementary Text**. The mathematical model yields motif occurrences that match those in random networks only when correlations are absent (i.e., *λ* = 0; Equations S2-3 in the **Supplementary Text**). In turn, for statistical ensembles of connectomes with correlations, the mathematical model allows us to explore how sparsity and heterogeneity in connectivity can in principle affect motif occurrences. For example, in sparsely connected networks (e.g. mean connection probability of 10%) feedforward motifs become increasingly underrepresented with increasing heterogeneity **(Fig. 4C)**. Conversely, in densely connected networks (e.g. mean of 90%) such motifs become increasingly overrepresented with increasing heterogeneity. Recurrent loops are always overrepresented in the presence of correlations, and overrepresentation increases the sparser and the more heterogeneous a network is interconnected **(Fig. 4C)**.

We test whether correlations, in conjunction with sparsity and heterogeneity, could indeed explain our observation that the structural composition of the neuropil results in nonrandom network architectures. We illustrate these quantifications for excitatory neurons in layer 5 **(Fig. 5A)**. About 80% of these neurons represent either intratelencephalic (L5ITs), pyramidal tract (L5PTs) or corticocortical neurons (L6CCs) **(Fig. 5B)**. We therefore analyze the pairwise statistics with respect to these morphological cell types by grouping the connection probability values in the statistical ensemble of connectomes accordingly **(Fig. 5C)**. Even though neuron somata from all of these cell types intermingle, the shapes of the respective connection probability distributions differ substantially. For example, L5ITs are predicted to connect more densely and less heterogeneously to L5PTs than L5PTs connect to one another **(Fig. 5D)**. Within the population of L5PTs, the shapes of connection probability distributions differ substantially depending on how far apart their somata are **(Fig. 5D)**. Thus, in any network from the statistical ensemble of connectomes, the degrees of sparsity and heterogeneity in connectivity will reflect the locations of neurons and their respective morphological properties **(Fig. S4A)**.

**Figure 5.**
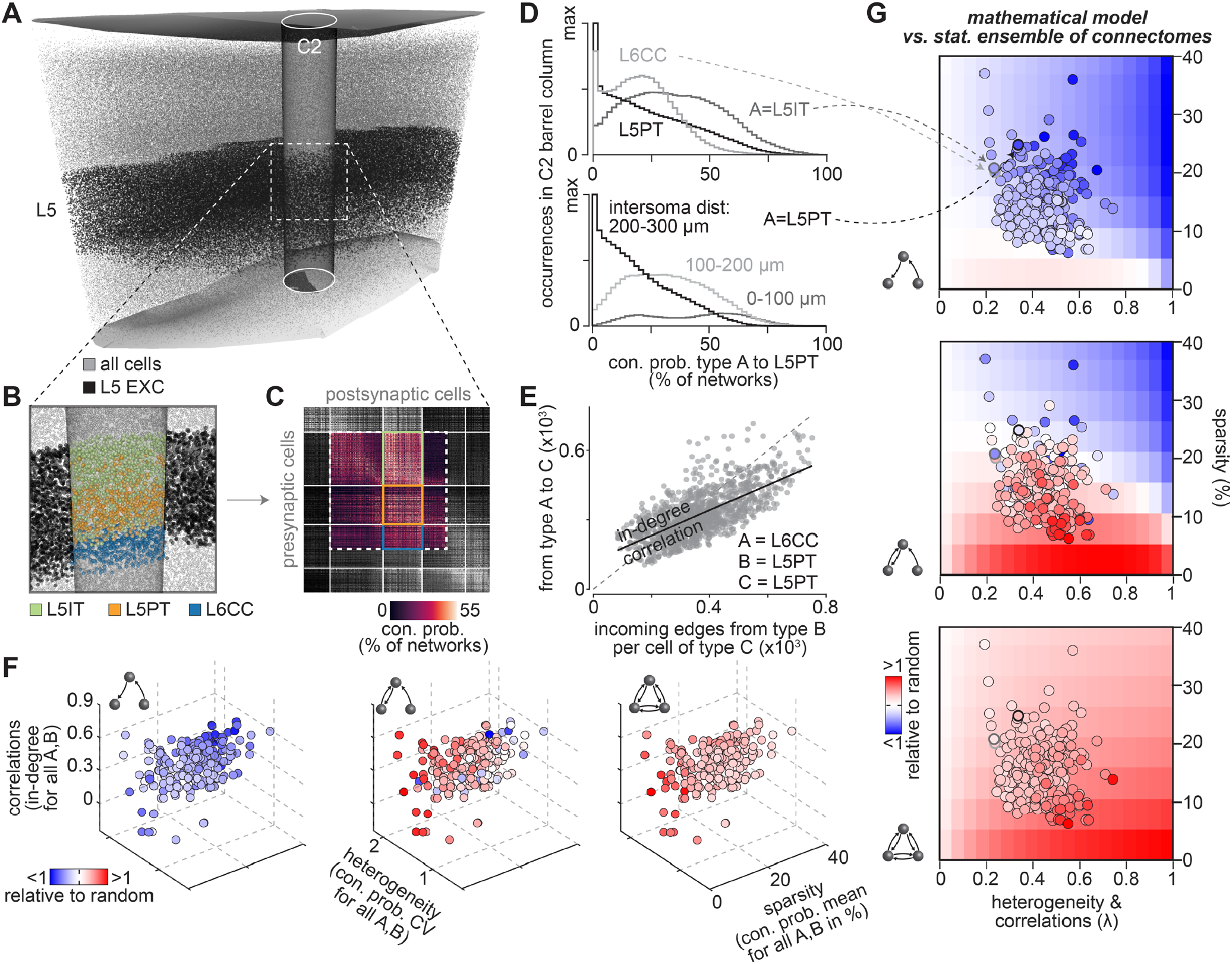
Three morphology-related factors define nonrandom network topologies. **(A)** The impact of neuron morphology on network architecture is illustrated for excitatory neurons in layer 5 (L5 EXC) of the C2 barrel column. **(B)** Zoom-in shows somata colored by their respective cell type. **(C)** Part of the matrix representation of the statistical ensemble of connectomes from **Fig. 1E** that represents the neurons shown in panel B. **(D)** Distributions of connection probabilities from the matrix in panel C for different cell type combinations. Bottom panel: connection probability distributions between L5PTs for different inter-somatic distances. **(E)** In-degree distributions derived from the matrix in panel C. The numbers of incoming connections that L5PTs receive from one another correlates with those they receive from L6CCs. **(F)** Motif occurrences in networks from the statistical ensemble of connectomes versus those in random networks depend on the in-degree correlation coefficients, the means and CVs of the corresponding connection probability distributions. Circles represent more than 200 groupings that represent neurons with different cell type combinations (e.g. those in panel D). **(G)** Nonrandom motif occurrences for the groupings in panel F are consistent with the mathematical model.

We find substantial correlations in connectivity within the statistical ensemble of connectomes. For example, L5PTs are predicted to receive more connections from one another the more connections they receive from L6CCs **(Fig. 5E)**. These ‘in-degree’ correlations reflect similarities in axon projection patterns. More specifically, despite substantial quantitative and qualitative morphological differences between L5PTs and L6CCs, the axons of both populations predominantly innervate the deep layers, where they span horizontally across several barrel columns (*26*). In contrast, L5IT axons predominately innervate the upper layers and remain largely confined to the dimensions of a single barrel column in the deep layers. As a result, in-degree correlations are much weaker between L5ITs and the other two cell types **(Fig. S4B)**. Thus, in any network from the statistical ensemble of connectomes, correlations in connectivity will be present, and the strengths of these correlations will reflect how similar the neurons’ locations and their respective morphologies are **(Fig. S4C)**.

We show that these observations generalize to more than 200 groupings that represent neuron populations of different cell type combinations **(Fig. 5F)**. For any of the groupings, motif occurrences deviate from those in random networks. The sparser and the more heterogeneous neurons are interconnected within a particular group, the more overrepresented are recurrent connections between them. Conversely, the more densely the group is interconnected, the more underrepresented are feedforward connections. These relationships are consistent with the mathematical model **(Fig. 5G)**. Thus, the shapes and correlations of pairwise connectivity statistics that reflect the structural composition of the neuropil represent a defining source for nonrandom network architectures. Consequently, the specific topology of cortical networks will reflect the specific morphological properties of their constituents.

### Principles linking neuron morphology to network architecture

The statistical ensemble of connectomes reveals four principles by which the neurons’ morphologies can impact network architectures. First, the vast majority of neurons whose axons and dendrites overlap remain unconnected. As a result, the more neuronal processes are packed into the neuropil relative to the number of synaptic structures that they represent, the smaller the probability that their respective pre- and postsynaptic structures could be connected to one another. Thus, the packing density of the neuropil translates into the means of connection probability distributions, and thereby defines a networks’ sparsity **(Fig. 6A)**. Second, the higher the diversity of neuronal processes that overlap (i.e., with respect to the morphologies of the neurons from which these processes originate) the broader are the distributions of connection probabilities. Thus, morphological diversity translates into the CV of connection probability distributions, and thereby defines a network’s heterogeneity **(Fig. 6B)**. Third, the more similar the dendrite or axon projection patterns of neurons are, the more similar are their respective contributions to the structural composition of the neuropil across subvolumes. Thus, similarities in the neurons’ projection patterns translate into correlations between connection probability and degree distributions **(Fig. 6C)**. Fourth, due to these correlations, the degrees of sparsity and heterogeneity define a network’s specific nonrandom topology. High packing density and high morphological diversity, as for example in the cerebral cortex, thereby yield recurrent network architectures **(Fig. 6D)**. Tissue with low packing density and low diversity yields feedforward architectures.

**Figure 6.**
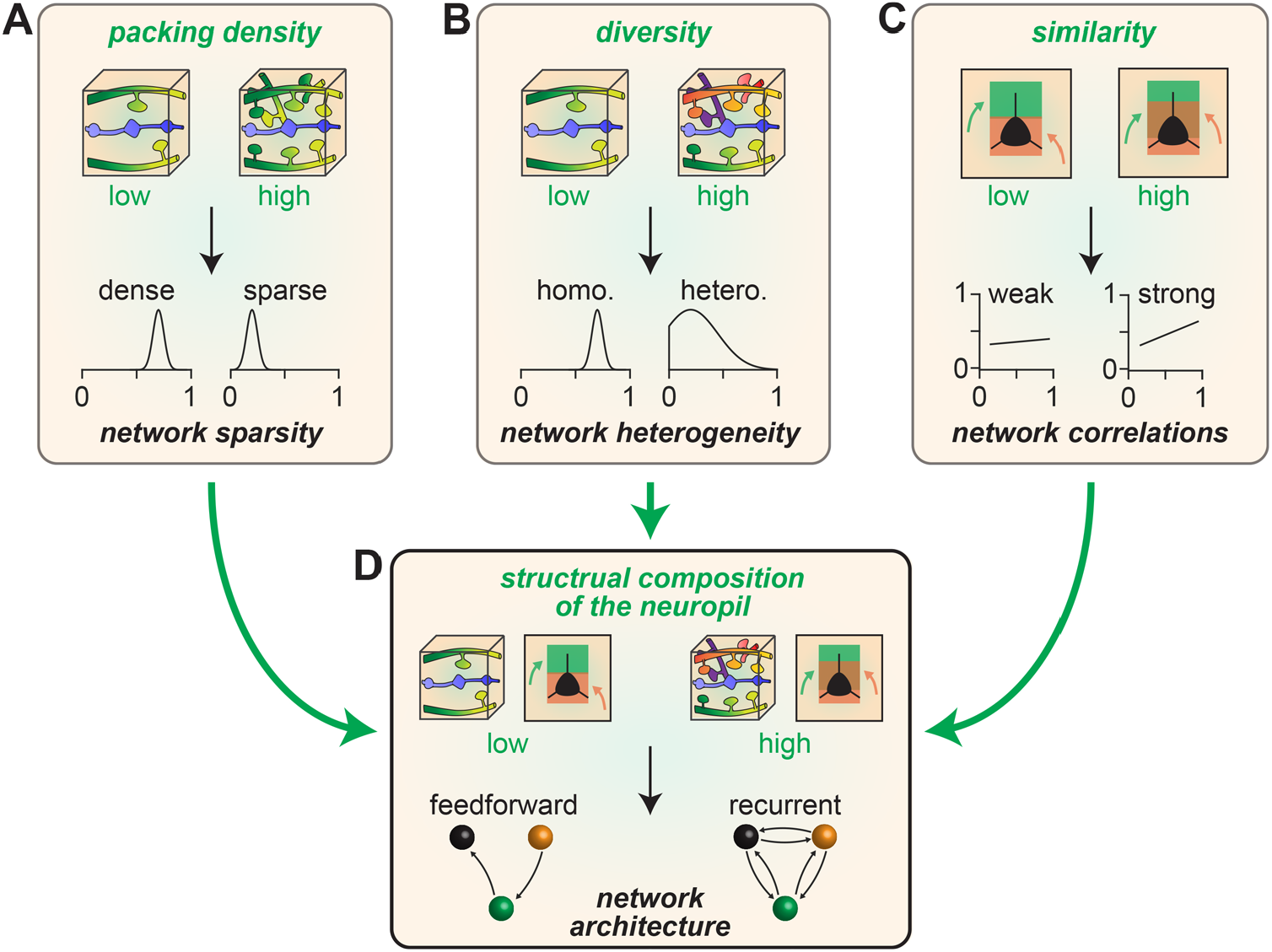
Principles linking neuron morphology to network architecture. **(A)** The packing density of neuronal processes within the neuropil translates into the means of connection probability distributions, and thereby defines a networks’ sparsity. **(B)** The diversity of neuronal processes that overlap (e.g. with respect to the morphologies of the neurons from which these processes originate) translates into the widths of connection probability distributions, and thereby defines a network’s heterogeneity. **(C)** Similarities in dendrite and axon projection patterns translate into correlations of connection probability and degree distributions. **(D)** In the presence of correlations, the degrees of sparsity and heterogeneity define a network’s specific nonrandom topology. Feedforward motifs become increasingly overrepresented the denser and the more homogeneously neurons are interconnected (i.e., low packing density and low morphological diversity). Conversely, recurrent motifs become increasingly overrepresented the sparser and the more heterogeneously neurons are interconnected.

### Structural composition of the neuropil could underlie empirically observed wiring specificity

How strong is the impact of neuron morphology on network architecture? Here we can address this question, because the statistical ensemble of connectomes provides a null hypothesis for testing to what degree empirically observed wiring specificity could reflect the structural composition of the neuropil, or whether it must be due to specific synapse formation mechanisms. We compare the predictions of the statistical ensemble of connectomes with a rich body of literature that represents several decades of quantitative connectivity measurements by different laboratories. First, we test whether the predicted relationship between packing density and sparsity in connectivity is observed empirically. We generate 500,000 µm^3^ subvolumes in layer 4 of the structural model **(Fig. S1C)** analogous to dense reconstructions in mouse barrel cortex (*13*). On average, these subvolumes comprise more than 10^8^ axon-dendrite pairs. In any network from the statistical ensemble of connectomes, the vast majority of these branch pairs remain unconnected **(Fig. 7A)**, while ∼10^5^ are predicted to be connected by a single synapse, ∼4,000 by two synapses, ∼150 by three synapses and ∼20 by four or more synapses **(Fig. 7B)**. These predictions for both the packing density **(Fig. S1D)** and the resulting degrees of sparsity are remarkably consistent with the empirical observations in mouse barrel cortex (*13*).

**Figure 7.**
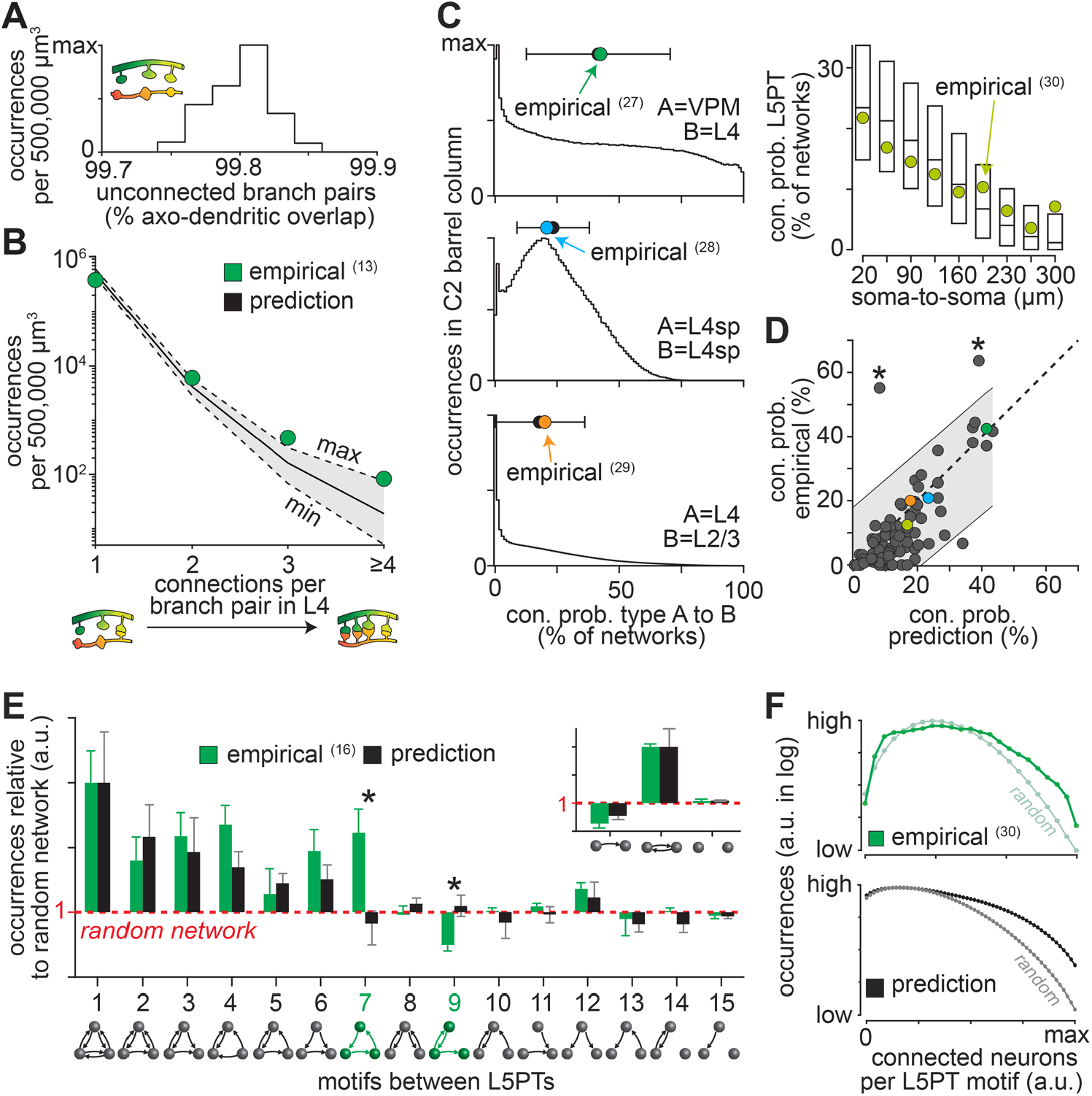
Predicted versus empirical connectivity data. **(A)** Predicted occurrences of unconnected branch pairs per 500,000 µm^3^ large subvolumes in layer 4 of the structural model of rat barrel cortex (n=252). **(B)** Occurrences of branch pairs connected by one or more synapses for the same subvolumes as in panel A match with empirical data from mouse barrel cortex (*13*). **(C)** Predicted connection probabilities match those of four example studies (*27-30*). **(D)** Empirical connection probabilities for 89 layer and/or cell type groupings versus those predicted here **(Tables S1-3)**. Grey shading represents 95% prediction interval. Asterisks denote inconsistencies with empirical data. **(E)** Predicted nonrandom motif occurrences between L5PTs match with the empirical data (*16*). As reported by (*16*), we normalized the triplet ratios by the predicted occurrences of doublet motifs (inset). Y-axis in log scale. **(F)** Overrepresentation of motifs between eight L5PTs increases with the number of connected neurons empirically (*30*) and in the statistical ensemble of connectomes.

Second, we test whether the predicted relationship between morphological diversity and heterogeneity in connectivity is observed empirically. For this purpose, we group neurons analogous to measurements that sampled connectivity between neuron pairs depending on their soma locations within a particular layer, cell types, inter-somatic distances, or combinations thereof **(Fig. 7C)**. In total, we predict the pairwise connectivity for eighty-nine such samplings that reflect different locations and/or morphological properties, and compare those with the respective empirical data reported across a set of twenty-nine studies **(Tables S1-3)**. The predicted connection probabilities correlate significantly with the empirical data (R=0.75, p<10^−16^). About 2/3 of the empirical connectivity values deviate from the prediction by less than half a standard deviation of the respective connection probability distribution, 94% by less than one standard deviation **(Fig. 7D)**. Random permutations of the predicted connection probabilities yield correlations with the empirical data that are not significant (R=0.00 ± 0.11).

Finally, we test whether the predicted nonrandom occurrences of motifs are observed empirically. The occurrences of all fifteen triplet motifs and their respective deviations from a random network were systematically assessed for L5PTs (*16*). The statistical ensemble of connectomes predicts nonrandom motif occurrences for this cell type that are remarkably consistent with these empirical data, with the notable exception of the feedforward loop and the recurrent feedback motifs **(Fig. 7E)**. Moreover, probing the occurrences of motifs between up to eight L5PTs revealed that independent of their particular topology, motifs become increasingly overrepresented with increasing numbers of connected neurons (*30*). This relationship is qualitatively consistent with our predictions **(Fig. 7F)**. Thus, wiring specificity from subcellular to network scales that was observed empirically between excitatory neurons – i.e., the occurrences of clusters of synapses; layer-, cell type- and distance-specific connectivity; over- or underrepresentation of motifs – could reflect the specific structural composition of the neuropil.

### Outlook: Disentangling sources of wiring specificity in densely reconstructed cortical networks

Our findings will advance the analysis and interpretation of future connectomics data – i.e., as soon as dense electron microscopy reconstructions become available for large cortical volumes. To illustrate how such datasets could be analyzed in principle, we apply our statistical approach to a densely reconstructed volume that comprises one cubic millimeter of human cortex (*23*). We analyze this petascale dataset in the preliminary form in which it was reported, which however reflects the current state-of-the-art for dense reconstructions of cortical tissue. We divide the dataset into small subvolumes and split all synaptic connections therein into pre- and postsynaptic structures along the reconstructed axonal and dendritic processes **(Fig. 8A)**. Analogous to the structural model of the barrel cortex, we then derive all network configurations that could emerge from the hence quantified structural composition of the neuropil, and calculate how likely these networks are to appear in the absence of synapse formation mechanisms that introduce wiring specificity **(Fig. 8B)**. The resulting statistical ensemble of connectomes for this fragment of human cortex predicts nonrandom network architectures that are remarkably consistent with the current, yet incomplete, stage of the automated reconstruction **(Fig. 8C)**. Interestingly, the predicted occurrences of the feedback motif deviate from the reconstruction for some groupings of pyramidal neurons **(Fig. 8D**). These preliminary results indicate that our statistical approach sets the stage to quantitatively disentangle whether wiring specificity in dense reconstructions is due to specific synapse formation mechanisms (e.g. occurrences of feedback motifs between particular pyramidal neurons), or whether it could reflect the structural composition of the neuropil (e.g. occurrences of other motifs between these neurons; **Fig. 8D**).

**Figure 8.**
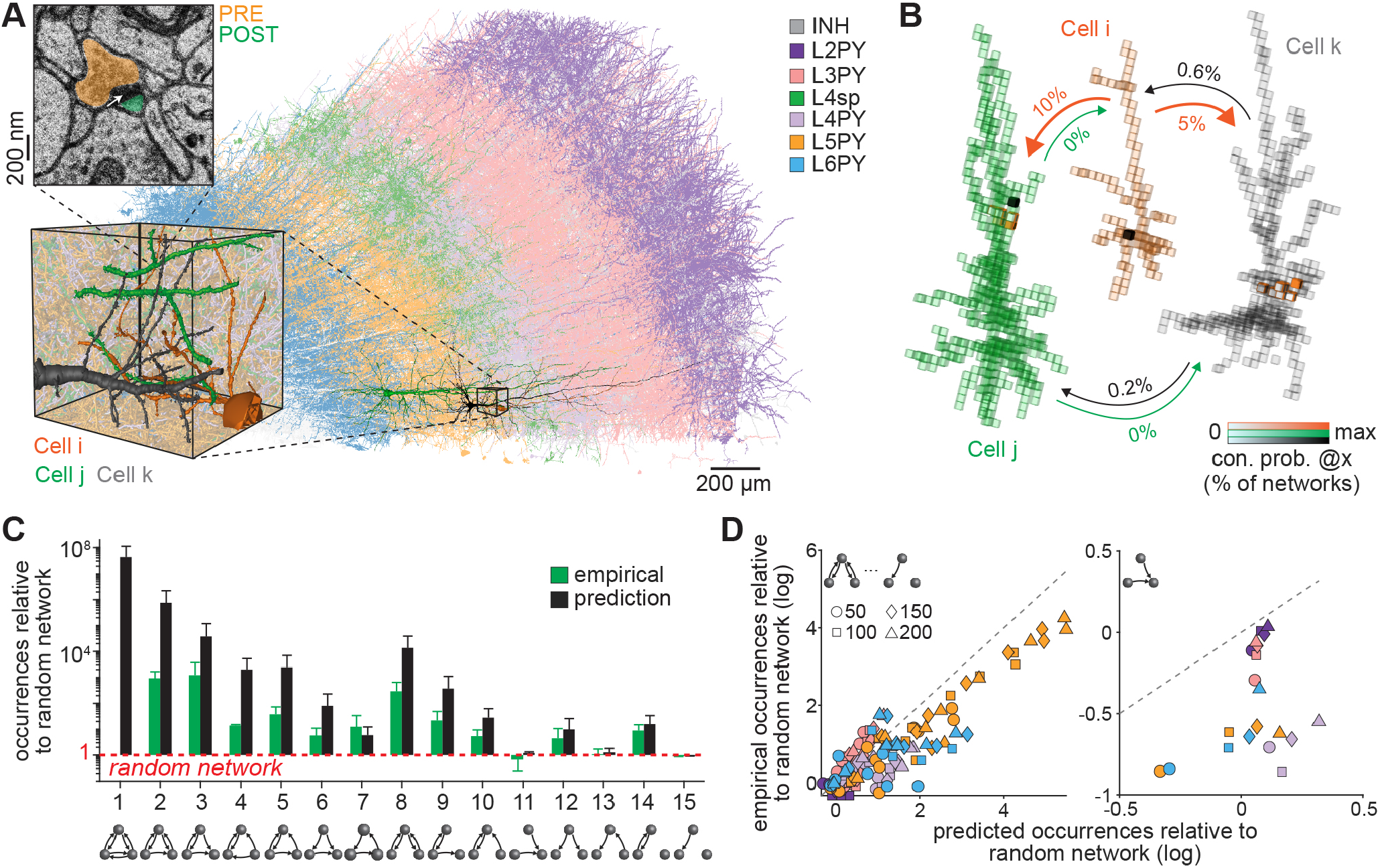
Testing the impact of neuron morphology on network architecture via dense reconstructions. **(A)** Current stage of the dense reconstruction of one cubic millimeter of human cortex, as recently reported (*23*). We colored the automatically reconstructed neuron morphologies by their laminar soma locations and putative morphological cell types (analogous to **Fig. S1**). Zoom-in shows three examples of layer 5 pyramidal neurons (L5PY). The electron microscopy image shows one synapse between the example neurons. We split all reconstructed synapses in this dataset into presynaptic (i.e., boutons) and postsynaptic structures (e.g. spines). **(B)** Analogous to **Fig. 1**, we divide the dense dataset into small subvolumes, count the pre- and postsynaptic structures along axons and dendrites therein, and generate the different network configurations which can account for them. We illustrate the resulting statistical ensemble of connectomes for the example neurons from panel A. Arrows denote the likelihoods that e.g. *Cell i* is connected by at least one synapse to the dendrites of *Cell j* within one of their overlap volumes (i.e., in 10% of the network configurations). **(C)** Predicted nonrandom motif occurrences between the 200 most connected neurons per layer match with the preliminary reconstructions. Error bars represent standard deviations across layers. Note that motif 1 has yet not been observed in the reconstructions of the inspected samples. **(D)** Predicted nonrandom occurrences of triplet motifs for the 50, 100, 150, and 200 most connected pyramidal neurons per layer are consistent with the preliminary reconstructions (left), except for the feedback motif (right).

## Discussion

We introduce the concept of statistical ensembles of connectomes to quantitatively test the impact of neuron morphology on cortical network architecture. We apply this strategy to an anatomically detailed structural model of the rat barrel cortex, which we show provides realistic and robust estimates for the structural composition of the neuropil for this entire brain area. We show that all network configurations that can emerge from the structural model are neither random nor consistent with Peters’ rule – two observations so far considered to be evidence for synapse formation mechanisms that connect neurons depending on their activity or genetically defined cellular identity. We find that these nonrandom properties reflect three factors of the structural composition of the neuropil that translate into a network’s sparsity, heterogeneity, correlations and topology. Thus, we demonstrate that in contrast to Peters’ rule, the impact of neuron morphology on network architecture cannot be direct in that it predicts connectivity. Instead, neuron morphology indirectly impacts network architecture by shaping the structural composition of the neuropil – i.e., the specific properties of the neuropil translate into specific pairwise and higher-order connectivity patterns that emerge irrespective of which particular axons and dendrites that share the same subvolume are connected.

We emphasize that our findings do not imply that the impact of specific synapse formation on wiring specificity is negligible (e.g. motifs 7 and 9 in **Fig. 7E**; motif 11 in **Fig. 8D**). Especially for inhibitory neurons, such a conclusion is not justified. Depending on their cell type, axons of inhibitory neurons preferentially target specific cell types and specific cellular compartments (reviewed in (*31*)). Testing the degrees to which the morphologies of inhibitory neurons impact network architectures would hence require incorporating the target specificity of different inhibitory cell types into the wiring rules that generate the statistical ensemble of connectomes. For now, however, relationships between morphology and target specificity of inhibitory neurons are not fully resolved. Thus, we limited our analyses to excitatory intracortical and thalamocortical connections for which empirical connectivity data is accessible. For these excitatory connections, we find that the structural composition of the neuropil predicts connectivity patterns that are in remarkable agreement with those observed empirically.

This consistency does not imply any causal relationship between neuron morphology and network architecture (i.e., axo-dendritic proximity does not predict connectivity). Instead, this consistency raises the question which synapse formation mechanisms could implement such strong relationships between network architectures and the neuropil’s structural composition. Here, the statistical ensemble of connectomes is based on assumptions that are consistent with nonspecific synapse formation mechanisms, for example, that axons compete with one another to connect to the available postsynaptic structures. Such mechanisms represent a prominent wiring strategy during the development of the peripheral (*32, 33*) and central nervous systems (*34*). It is hence tempting to speculate that neuron morphology development, in conjunction with competitive synapse formation mechanisms, establishes a scaffolding of wiring properties in the architectures of cortical networks. If this were true, the impact of neuron morphology on cortical network architecture might diminish throughout life. Consistency with the empirical data could thereby reflect the fact that connectivity measurements in the literature originated largely from young animals **(Table S1)**. This interpretation is supported by dense reconstructions of the nematode C. elegans, which recently revealed that the structural composition of its nervous system provides a constant scaffolding on which connectivity is remodeled from birth to adulthood (*35*). Interestingly, some structurally determined wiring patterns were maintained throughout life. This may also apply to the cerebral cortex, as its proper function is critically linked to two network properties that we found to be strongly impacted by morphology: heterogeneity and correlations in connectivity (*36*). Consequently, to ensure robustness of cortical dynamics, homeostatic mechanisms may maintain structurally determined wiring specificity despite the constant remodeling of cortical networks. As illustrated in **Fig. 8**, the concept of statistical ensembles of connectomes will allow testing whether the impact of morphology on cortical network architecture decreases or is maintained during maturation.

Scaffoldings of wiring properties that emerge in cortical networks during development were proposed to reflect an evolutionary strategy for implementing innate sensory representations and behaviors (*37*). However, what exactly genomes specify about wiring remains unknown. In C. elegans, the genome has the capacity to specify every connection between every neuron in minute detail. In contrast, cortical synapses cannot be specified so precisely, even if the entire genome would solely encode for connections. Due to this ‘genomic bottleneck’, it was suggested that scaffoldings in cortical networks are compressed in the genome as wiring rules (*37*). However, we suggest that the emergence of scaffoldings in networks does not rely on such explicitly encoded wiring rules. Instead, scaffoldings could be encoded implicitly via genetically induced developmental programs that guide neurons and their processes into specific subvolumes of the cortical sheet (*38*). Implicit encoding would not only solve the issue of compression through the genomic bottleneck. It may also explain how cortical networks adapt to the environment. For example, periphery-driven activity can regulate guidance programs that shape neuron morphologies (*39*). Accordingly, sensory experience could alter how the structural composition of the neuropil impacts cortical network architecture. Implicit encoding of wiring properties via neuron morphology development may hence reflect an efficient evolutionary strategy to maintain cortical network architectures across generations while providing sufficient flexibility for invading new ecological niches (*40*).

## Materials and Methods

All relevant data and codes are available from the authors. We used custom-written routines in C++, Python, or MATLAB 2020b software (Mathworks, Natick, MA, USA) for analysis and Amira software for visualization. Boxplots were generated with the Matlab built-ins *boxplot* or *boxchart* where the bottom and top of the box represents the 25^th^ and 75^th^ percentiles, and the line within the box the median. The lines extend to the adjacent values. Outliers are all values more than 1.5 times the interquartile range away from the top or bottom of the box.

### Structural model of rat barrel cortex

#### NeuroNet

We reverse engineered the structural composition of the neuropil for the rat barrel cortex by using NeuroNet, a custom-designed extension package for Amira software (FEI). NeuroNet was described in detail previously (*19*). Briefly, NeuroNet requires the following anatomical data as input: (i) a reconstruction of the 3D geometry and cytoarchitecture for the cortical volume of interest, (ii) a spatially dense reconstruction of the distributions of excitatory (EXC) and inhibitory (INH) neuron somata within the volume, (iii) samples of *in vivo* labeled dendrite and axon reconstructions that represent neurons from all layers and for all major morphological cell types, and (iv) cell type- and target layer-specific measurements for the densities of pre- and postsynaptic structures along these axons and dendrites, respectively. The output by NeuroNet is a digital model of the cortical volume, where each neuron soma is represented by one axon and dendrite from the sample of morphologies **(Fig. S1)**.

#### Anatomical input data

All anatomical data used here as input for NeuroNet was acquired in Wistar rats (primarily during the fifth postnatal week) and has been reported previously. Briefly, to capture the characteristic geometrical, cytoarchitectonic, and cellular organization of the rat barrel cortex in the structural model, we reconstructed precise 3D maps of cortical barrel columns with surface reconstructions of the pia and white matter (*41*), and quantified the locations of all EXC and INH neuron somata in the rat barrel cortex and in VPM thalamus (*42*). To capture the cell type-specific morphological organization of the rat barrel cortex in the structural model, we reconstructed a sample of *in vivo* labeled EXC neuron morphologies (*26, 43, 44*) and the intracortical part of *in vivo* labeled VPM axon morphologies (*45*). NeuroNet replaced each neuron of the reconstructed distribution of EXC somata with a morphology from this sample of reconstructions. The neurons’ locations in the structural model were on average within ±119 µm of their reconstructed 3D soma positions (*41*). We placed as many thalamocortical axons as the average measured number of neurons per respective VPM barreloid. To account for EXC connections onto INH neurons, we incorporated reconstructions of INH neurons into the structural model (*46-49*). For *in vitro* labeled INH neurons, morphologies were extrapolated by assuming radial symmetry. Connections from or onto INH neurons were hence not systematically analyzed. To capture the rat barrel cortex’ distribution of synaptic structures in the structural model, we derived the number of presynaptic structures (i.e., axonal boutons) by multiplying the axon length that each neuron contributes to a particular subvolume with the number of boutons per length (*19*), as measured for all EXC cell types and layer 1 INH neurons, and depending on the axons’ target layer (*26, 49*). For all remaining INH neurons we set the density to 0.2 boutons per µm axon as reported in (*50-52*). Based on the resulting density distribution of boutons along the cortical depth, we scaled the total number of postsynaptic structures along the dendrites. More specifically, we performed the scaling separately for the targets of boutons from EXC and INH axons: First, for targets of EXC boutons, we derived the number of postsynaptic structures along EXC dendrites (i.e., spines) by assuming that spine densities are proportional to dendritic length. For the respective number of postsynaptic structures along INH dendrites and somata, we assumed proportionality to their respective surface areas. The derived density of postsynaptic structures for EXC neurons ranged from 1.04 to 1.68 spines per µm dendritic length, consistent with empirical spine density measurements (*53, 54*). The derived density of postsynaptic structures for INH neurons was 0.74 per µm^2^ of dendritic or somatic surface, consistent with empirical synapse density measurements on INH somata (*55, 56*). Second, for targets of INH boutons, we derived the number of postsynaptic structures along both EXC and INH dendrites and somata by assuming proportionality to their respective surface areas. The derived density of postsynaptic structures for EXC and INH neurons was 0.06 per µm^2^ of dendritic and somatic surface, consistent with empirical data (*55-57*).

#### Validation of the structural model

We used NeuroNet to create >30,000 structural models of the barrel cortex with different anatomical data as input and quantified the variability across structural models for 512 (50 µm)^3^ large subvolumes within layers 2 to 6 of the C2 barrel column – i.e., axonal and dendritic path length, number of branches, number of branches that remain unconnected to a soma within the same subvolume, number of synaptic structures (e.g. boutons), number of contributing cells and cell types, and path length to the soma from each branch **(Fig. S1D/E)**. First, we assessed how these parameters of the structural composition are affected by the limited sample size of morphology reconstructions. We generated structural models where only one morphology per cell type, two morphologies, and so on were used as input to NeuroNet. Starting with one morphology per type, we used a random sample of 500 combinations of morphologies as input to generate 500 structural models. All structural models were based on the same distribution of neuron somata (i.e., average across four barrel cortices (*42*)). For each of the subvolumes, we determined the CV of each structural feature across the 500 structural models. We then calculated the median CV of each structural feature across all subvolumes. We refer to the median CV as the ‘morphological uncertainty’ per subvolume **(Fig. S1F)**. We repeated this analysis for two morphologies per cell type and so on until the maximal sample size of morphologies was reached, respectively. Second, we assessed how the parameters of the structural composition are affected by the variability of soma distributions across animals. For this purpose, we repeated the analysis of ‘morphological uncertainty’ with structural models that were based on each of the four measured dense distribution of neuron somata that were used to create an average distribution (*42*). We again calculated the median CV of each structural feature across all subvolumes and refer to the median CV as the ‘cellular uncertainty’ per subvolume **(Fig. S1F)**.

#### Statistical ensemble of connectomes

We derived the statistical ensemble of connectomes for the structural model of rat barrel cortex **(Fig. S1)** by assuming that only the presence of pre- and postsynaptic structure within the same subvolume is necessary for synapse formation – i.e., their particular positions or proximity (e.g. touch) within a subvolume are not taken into account (*19*). First, we calculated the *dense structural overlap* (***DSO***) as the product of the numbers of pre- and postsynaptic structures that neurons ***i*** and ***j*** contribute to a subvolume ***x***, relative to the total number of postsynaptic structures contributed by all neurons, here indexed with ***N***.

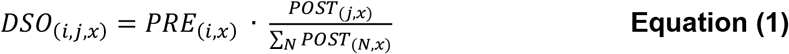

Based on this quantity, we assume that any presynaptic structure has equal probability of forming a connection with any of the allowed postsynaptic structures present within the same subvolume. The probability ***p*** for the presence of ***n*** connections between neurons ***i*** and ***j*** within a subvolume ***x*** across all networks is therefore given by a Poisson distribution with parameter ***n***:

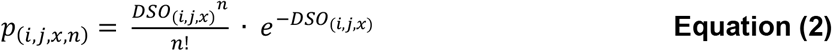

We assume that the formation of connections does not affect synapse formation elsewhere. Thus, the probability ***P*** that neurons ***i*** and ***j*** are connected by ≥1 synapse across all networks is given by:

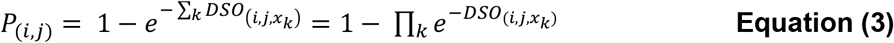

where the index ***k*** runs over all subvolumes in which neurons ***i*** and ***j*** overlap. Parameterizing the subvolumes of the structural model of the barrel cortex by the quantity ***DSO***, followed by application of Equations 2-3, yielded the statistical ensemble of connectomes that we analyzed here. We analyzed the statistical ensemble of connectomes for subvolumes of (50 µm)^3^ unless stated otherwise (e.g. **Fig. 3**). The color map **Figure 1E** represents 0-95% of the connection probability values (i.e., 0-55%).

### Quantification and statistical analysis

#### Testing of Peters’ rule

To test the validity of Peters’ rule at the subcellular level we restricted our analysis to layers 2 to 6 within the C2 barrel column. Within that volume, we calculated the number of boutons and branch pairs that form zero to eight or more synapses per (50 µm)^3^ large subvolumes (n=512). We repeated this analysis for 512 subvolumes with 1, 5, 10, and 25 µm edge lengths and for 64 subvolumes with 100 µm edge length. At 1 µm edge length we excluded 229 subvolumes where either no axon or no dendrite were present. To test Peters’ rule at the cellular level we restricted our analysis to a combination of ∼400 million neuron pairs. For each pair, we determined the number of (50 µm)^3^ large subvolumes with axo-dendritic overlap, referred to as ***n***_***overlap***_. We then calculated in how many networks the pair forms zero to ***n***_***overlap***_ synapses across all subvolumes. The resulting occurrences represent an upper bound since we constrained the overall number of connections and not the number of connections per subvolume. We mapped the number of connections per pair on bins of 1% width ranging from 0% (no connection between pair) to 100% (as many connections as ***n***_***overlap***_). Finally, we determined the average number of occurrences across all pairs per bin. The resulting profile was smoothed with a moving median for visualization purposes. We determined how many of ∼400 million pairs overlap at 1, 5, 10, 25 and 100 µm edge length and how often those were connected in the structural model of the barrel cortex.

### Analyses of statistical ensemble of connectomes

We used the Matlab built-in *digraph* to illustrate network configurations for 50 neurons as a graph. Edges between neurons were realized based on their predicted connection probability in the statistical ensemble of connectomes. We constrained each configuration to have the same number of edges. We generated the random network example by randomly assigning the same number of edges to 50 neurons. To analyze network topologies, we calculated the occurrence probability of each of the 15 motifs for a set of 8 million randomly selected neuron triplets with each neuron belonging to a particular neuron population (e.g. neurons were grouped by their cell type or soma position in a layer). We calculated the mean probability of the occurrences for each motif as predicted by the statistical ensemble of connectomes and compared it with those expected in a random network: First, we calculated the mean connection probabilities for each of the six edges between all of the sampled neurons. Second, we used these six mean connection probabilities to calculate the occurrence probability of each motif. Third, we divided the predicted motif probabilities by their respective expected probabilities in the random network. We calculated the deviation of motif occurrences of all 15 motifs for all 220 cell type-specific triplet combinations. For each triplet combination we calculated the mean and CV of their connection probability distribution across all six edges. For triplets with neurons from at least two different cell types (n=210), we also calculated the mean across all in-degree correlation coefficients involving these cell types. We extended our analysis to motifs between more than three neurons. We computed the probabilities of motif 1 (recurrent loop) and 13 (feedforward chain) for up to ten neurons. For this purpose, we randomly sampled sets of neurons per motif size (10 million for each motif) from the statistical ensemble of connectomes (i.e., 3 to 10) and computed the occurrence probabilities, respectively. We then compared the respective probabilities with those computed in random networks based on the mean connection probability across all neurons of the sample. We computed the mean, standard deviation, and CV, of the connection probabilities between all groupings. We assessed correlations between neurons by calculating the number of incoming edges ***n***_***EDGES***_ between presynaptic neurons ***i*** (including those from VPM) and postsynaptic neurons ***j*** by summing their respective ***DSO*** across all overlapping subvolumes ***x***:

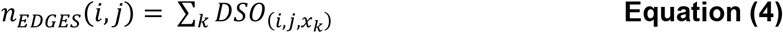

We grouped all pre- and postsynaptic neurons by their cell type identity, and summed ***n***_***EDGES***_(***i, j***) across each presynaptic population. This resulted in the mean number of connections (i.e., in-degree) each postsynaptic neuron ***j*** receives from this population across networks. We computed a linear regression fit and Pearson’s linear correlation coefficient between the in-degrees of two different presynaptic populations onto all neurons per postsynaptic cell type. We repeated this computation for all combinations of presynaptic cell types.

#### Comparison with empirical connectivity data

We calculated the occurrences that two branches form zero, one, two, three, and (at least) four synapses for all branch pairs for a sample of 252 subvolumes of 100×100×50 µm^3^ in layer 4 for comparison with the empirical data reported for layer 4 of mouse barrel cortex (*13*). We compared our predictions with 89 empirical connection probability measurements reported across a set of 29 studies (*11, 16, 27-30, 58-80*). We emulated the respective experimental conditions in the structural model. For empirical data from *in vitro* studies, we created twenty virtual brain slices of 300 µm thickness through the structural model. The slices were shifted by 20 µm with respect to one another along the rostro-caudal axis. We truncated the neuronal processes of all neurons whose somata were located within each slice (i.e., we cut branches at their intersection with the slice surface, and removed those branches from the structural model that became disconnected from the soma). We computed the connection probabilities between each neuron pair in the virtual slices as defined by **Equations 1-3** with the quantity ***DSO*** being the contribution of pre- and postsynaptic structures by the truncated pairs’ morphologies with respect to the total number of postsynaptic structures contributed by all neurons. We grouped the neurons as described in the respective studies (**Table S1**); i.e., by their laminar soma location and – if reported – by their cell type. Layer borders were defined as reported previously (*42*). If the recording depth underneath the slice surface was not reported, we restricted the comparison to pairs within the mean reported range of recording depths (31 µm to 130 µm). We computed the Pearson’s linear correlation coefficient between the empirical and predicted connection probabilities and the 95% confidence bounds for new observations based on a linear regression with no intercept using the Matlab built-ins *fitlm* and *predict*. We performed a random permutation test on the correlation coefficient by shuffling the empirical and the predicted connection probabilities and re-computing their correlation coefficient. We repeated this step 100,000 times. We compared the predictions with two empirical studies that performed connectivity measurements as a function of inter-somatic distance (*30, 58*) (**Tables S2-3**). Here, we grouped neurons additionally by their inter-somatic distance along the lateral axis (i.e., the axis running parallel to the slicing surface). We compared the predicted deviations of motif occurrences across L5PT triplets and doublets with empirical observations (*16*). We grouped the neurons accordingly and calculated the motif occurrences and ratios for ∼1.7 million L5PT doublets and ∼200,000 L5PT triplets across 20 slices through the structural model. We used the same analysis as reported by (*16*) and normalized the resulting triplet ratios by the doublet motif occurrences to avoid over- or underrepresentation of triplet motifs due to over- or underrepresentation of doublet motifs. We compared the motif probabilities across the number of edges in motifs of eight neurons to empirical observations (*30*). For this purpose, we randomly sampled 20,000 sets of eight L5PTs across the 20 slices through the structural model. For each set of neurons and each number of edges (ranging from 0 to 56 edges), we computed the number of edge combinations (e.g., 1 combination is possible for 0 or 56 edges, but more than 10^10^ combinations are possible of 10 edges). If the number of edge combinations was less than 1,000, we iterated over all combinations. If the number of combinations was larger than 1,000, we randomly generated 1,000 motifs that matched the number of edges. We calculated the respective (occurrence) probability of each edge motif in the slices through the structural model and a random network constrained by the respective mean connection probability.

#### Analysis of densely reconstructed volume from human cortex

We downloaded the annotated synapse dataset released with the C3 segmentation (gs://h01-release/data/20210601/c3/synapses/exported/) and filtered it for synapses whose presynaptic site was an axon and whose postsynaptic site was either a dendrite, soma, or axon initial segment, resulting in 133,704,943 synapses as described in (*23*). We divided the dataset into subvolumes of (25 µm)^3^ and assigned each synapse to a subvolume based on its location. Then we split each synapse into a pre- and postsynaptic structure. The dataset contains 15,567 reconstructions annotated as ‘neuron’. For each pair of neurons, we calculated the quantity *DSO* of its excitatory and inhibitory synapses separately. We then summed the excitatory and inhibitory *DSO* for each neuron pair over all subvolumes and computed the statistical ensemble of connectomes. We determined the 50, 100, 150, and 200 most connected neurons per layer. For each of these groupings, we determined how often each triplet motif occurred in the reconstructed dataset compared to a random network that has the same mean connection probability. We repeated this analysis for the statistical ensemble of connectomes as well as for samples of pyramidal neurons (annotated ‘pyramidal’).

## Acknowledgements

We thank Kevin Briggman, Peter L. Strick, Haim Sompolinsky, Idan Segev and David Fitzpatrick for discussions and comments on the manuscript. Funding was provided by the Center of Advanced European Studies and Research, the Max Planck Institute for Biological Cybernetics, Center for Neurogenomics and Cognitive Research, European Research Council under the European Union’s Horizon 2020 research and innovation program (grant agreement 633428; to M.O.), German Federal Ministry of Education and Research (grants BMBF/FKZ 01GQ1002 and 01IS18052; to M.O. and J.M.), Deutsche Forschungsgemeinschaft (SFB 1089 to M.O. and J.M.; and SPP 2041 to M.O., J.M. and H.H; and EXC number 2064/1, 39072764 to J.M.).

## Author contributions

M.O. conceived and designed the study. D.U. generated the data and developed the analysis. P.H. and H.H. developed analysis routines and created the online resource. J.M. developed the mathematical model. C.K. and B.S. provided data. D.U., P.H. and M.O. analyzed the data, and M.O. and D.U. wrote the paper with help from all authors.

