## Supplementary Material for "The Impact of Neuron Morphology on Cortical Network Architecture"

### Supplementary text

#### Structural model of rat barrel cortex

We had previously reverse engineered the structural composition of the neuropil for the volume of rat barrel cortex that represents the 24 major facial whiskers (*19*). This structural model provided quantitative estimates for the spatial distributions of all neurons, and their dendrites and axons – including the distributions of pre- and postsynaptic structures along these neuronal processes (reviewed in (*22*)). All anatomical data originated from rats of the same strain, sacrificed at similar time points following the critical periods of neuron morphology development. The structural model **(Fig. S1A)** comprises a cortical volume of 6.8 mm$^3$ (*41*), contains 477,551 excitatory and 69,788 inhibitory neurons, and is innervated by 6,225 neurons from VPM thalamus (*42*). These neurons give rise to 25.6 km of axonal and dendritic processes within the structural model, representing more than 5.5 billion pre- and postsynaptic structures, respectively **(Fig. S1B)**.

We validated that the structural model provides realistic estimates of the structural composition of a neuropil. This information is crucial for judging the validity of the derived statistical ensemble of connectomes. Consistent with early estimates (*81*), the structural model predicts that 3.3 km of axons and 0.5 km of dendrites are compressed into each cubic millimeter of cortical tissue. Such high packing densities represent a major challenge for dense electron microscopic reconstructions (*82*). The largest densely reconstructed cortical volume to date hence comprises only ~0.0005 mm$^3$ (*13*). This remarkable effort in layer 4 of mouse barrel cortex had provided first quantitative data about the structural composition of the cortical neuropil, and revealed that volumes with such dimensions **(Fig. S1C)** can comprise more than 45,000 neuronal processes, which represent ~2.7 m of path length and ~400,000 synaptic connections. These empirical observations fall well within the respective ranges predicted by the structural model **(Fig. S1D)**. Consequently, the configurations by which neuronal processes could be connected locally to one another in the structural model of the rat barrel cortex (i.e., within subvolumes <0.0005 mm$^3$) will be qualitatively and quantitatively very similar to those that could emerge from the densely reconstructed volume of mouse barrel cortex. Also consistent with the empirical data, the structural model predicts that >97% of the neuronal processes remain unconnected to a soma that lies within the same subvolume. The structural model thereby provides first insight into the locations and diversity of morphological cell types for the neurons from which these unconnected neuronal processes originate **(Fig. S1E)**.

The structural model predicted the number of neuronal processes and synaptic structures to vary substantially across the different subvolumes of the barrel cortex **(Fig. S1D/E)**. We therefore tested whether these variations of the structural composition are anatomically realistic. For this purpose, we generated >30,000 structural models of the rat barrel cortex, which reflected different measured soma distributions and/or samples of *in vivo* labeled dendrite and axon morphologies. This revealed that the structural composition of each subvolume would not change by more than 12% – the diversity of the morphological cell types from which neuronal processes originate by no more than 8% – even if the structural models were based on a larger sample of reconstructed morphologies **(Fig. S1F)**. Moreover, the structural models predicted layer-specific density variations of boutons **(Fig. S1G)** that are largely consistent with synapses density measurements in the barrel cortex of juvenile rats (*83*). Thus, the structural model analyzed here provides realistic and robust estimates for the amounts of axonal and dendritic branches within any subvolume of the rat barrel cortex, for the synaptic structures that these processes represent, and for the diversity of their respective cellular and cell type origins.

**Mathematical model for statistical ensembles of connectomes with correlations**

We consider our statistical ensemble of connectomes as a distribution of pairwise connection probabilities $\boldsymbol{p_i}$ that generates network configurations. Suppose that $\boldsymbol{K}$ connections are drawn from any such generating distribution $\boldsymbol{Q(p|\mu,\sigma)}$**,** where $\boldsymbol{\mu}$ and $\boldsymbol{\sigma}$ represent the mean (i.e., sparsity) and variance (i.e., heterogeneity) of the pairwise connectivity in the corresponding statistical ensemble of connectomes. If each of the $\boldsymbol{K}$ connections is drawn independently, the probability of observing for example recurrent loops (motif 1) is accordingly the expected value of $\boldsymbol{Q(p|\mu,\sigma)}$:

$$P(motif\ 1) = E_Q\left(\prod_{i=1}^{K} p_i\right) = \prod_{i=1}^{K} E_Q(p_i) = \mu^K \qquad \textbf{Equation (S1)}$$

Thus, when connection probabilities are independent of one another, motifs will occur as expected for randomly connected networks – i.e., occurrences are independent from the network's heterogeneity and only reflect the mean of the underlying pairwise statistics. Consequently, our observations of nonrandom occurrences of motifs, and their dependencies on network heterogeneity, cannot be consistent with the assumption that connection probabilities are independent of one another. Instead, only correlations in the statistical ensemble of connectomes could explain our observations. We therefore investigated how the presence of correlations affects the occurrences of motifs. For this purpose, we developed a mathematical model for correlated connectivity that is closely related to the one studied in (*25*). In the following, $\boldsymbol{\mathcal{N}(\mu,\lambda)}$ denotes a Gaussian distribution with mean $\boldsymbol{\mu}$ and variance $\boldsymbol{\lambda}$, $\boldsymbol{\varphi(t,\mu,\lambda)}$ denotes the respective Gaussian probability density function evaluated at $\boldsymbol{t}$, $\boldsymbol{\Phi(s,\mu,\lambda)}$ denotes the respective cumulative probability density function evaluated at $\boldsymbol{s}$, and the complementary cumulative probability density function is defined by $\boldsymbol{L(s,\mu,\lambda) = 1 - \Phi(s,\mu,\lambda)}$. For simplicity, the mathematical model assumes that whether there is an $\boldsymbol{i}$-th edge between two nodes (denoted by $\boldsymbol{X_i = 1}$, otherwise $\boldsymbol{X_i = 0}$) is the result of a combination of only one 'private' source $\boldsymbol{T_i}$, and one 'shared' source $\boldsymbol{S}$. The larger the shared source $\boldsymbol{S}$ is relative to the private one $\boldsymbol{T_i}$, the more correlated the resultant connection probabilities are. Here we found that such shared sources could originate from similarities in the neurons' locations and morphologies. The more similar the dendrite or axon projection patterns of neurons are, the more similar are their respective contributions to the structural composition of the neuropil across subvolumes, which leads to correlations between connection probability and degree distributions. The mathematical model could be easily generalized by incorporating more than one shared source (e.g. one for each cell type combination). The mathematical model has two parameters: $\boldsymbol{\lambda}$, bounded between 0 and 1, and representing the magnitude of the shared source, and thus the degree of correlation in the sources. As we will demonstrate in the following, $\boldsymbol{\lambda}$ also determines the heterogeneity of connection probabilities - the larger $\boldsymbol{\lambda}$ is, the more heterogeneous the connection probabilities are. The parameter $\boldsymbol{\gamma}$ represents the degree of connectivity – the greater $\boldsymbol{\gamma}$ is, the higher connection probabilities are. We define that the $\boldsymbol{i}$-th edge exists (i.e., $\boldsymbol{X_i = 1}$) whenever the joint input of $\boldsymbol{T_i}$ and $\boldsymbol{S}$, denoted by $\boldsymbol{Z_i}$, is larger than 0:

$$X_i = 1 \text{ whenever } Z_i > 0 \text{ where}$$

$$Z_i = \gamma + \sqrt{\lambda} S + \sqrt{\eta} T_i$$

where

$$\eta = 1 - \lambda\,,\ S \sim \mathcal{N}(0,1),\ T_i \sim \mathcal{N}(0,1)$$

$$Z_i \sim \mathcal{N}(\gamma, \eta + \lambda) = \mathcal{N}(\gamma, 1)$$

Thus, $\boldsymbol{cov(Z_i, Z_j) = \lambda}$. If $\boldsymbol{\lambda = 1}$, $\boldsymbol{X_i}$ is only determined by the shared source $\boldsymbol{S}$, while if $\boldsymbol{\lambda = 0}$, $\boldsymbol{X_i}$ is only determined by the private source $\boldsymbol{T_i}$. Given this mathematical model, the connection probability $\boldsymbol{p_i}$ for each edge $\boldsymbol{X_i}$ is given by:

$$p_i(S) = P(X_i = 1|S) = L(0, \gamma + \sqrt{\lambda} S, \eta)$$

And we likewise get

$$\begin{aligned}
\mu &= E_s(p_i) = L(0, \gamma, 1) \\
\sigma^2 &= Var_s(p_i) \\
&= \int_{-\infty}^{\infty} P(X_i = 1|s)^2 \varphi(s, 0,1) ds - \mu^2 \\
&= \int_{-\infty}^{\infty} L(0, \gamma + \sqrt{\lambda} s, \eta)^2 \varphi(s, 0,1) ds - \mu^2
\end{aligned}$$

Deriving the covariance between any two connections $\boldsymbol{X_i}$ and $\boldsymbol{X_j}$ yields:

$$\begin{aligned}
cov(X_i, X_j) &= E(X_i, X_j) - \mu^2 = P(X_i = X_j = 1) - \mu^2 \\
&= \int_{-\infty}^{\infty} P(X_i = 1|s) P(X_j = 1|s) \varphi(s, 0,1) ds - \mu^2 \\
&= \int_{-\infty}^{\infty} L(0, \gamma + \sqrt{\lambda} s, \eta)^2 \varphi(s, 0,1) ds - \mu^2 \\
&= Var(p_i)
\end{aligned}$$

Thus, in the simplified mathematical model the covariance of the connections is equal to the variance of connection probabilities – the more strongly the connection probabilities vary, the more strongly the connections themselves are correlated. Hence, the parameter $\boldsymbol{\lambda}$ represents a measure of both, the degree of correlation and heterogeneity. To assess the impact of $\boldsymbol{\lambda}$ and the mean connection probability μ onto motif occurrences and deviations, the probability that $\boldsymbol{k}$ out of $\boldsymbol{K}$ connections of a motif are realized is given by:

$$P(|X| = k) = \int_{-\infty}^{\infty} \binom{K}{k} L(0, \gamma + \sqrt{\lambda} s, \eta)^k \Phi(0, \gamma + \sqrt{\lambda} s, \eta)^{K-k} \varphi(s, 0,1) ds \qquad \textbf{Equation (S2)}$$

The probability of observing recurrent loops ($\boldsymbol{K = k}$) is accordingly:

$$P(motif\ 1) = \int_{-\infty}^{\infty} L\left(0, \gamma + \sqrt{\lambda}S, 1-\lambda\right)^K \phi(s,0,1)ds \xrightarrow{\lambda=0\ yields} L(0,\gamma,1)^K = \mu^K \quad \textbf{Equation (S3)}$$

This mathematical model was implemented as a numerical simulation in Matlab. We iterated over 250 $\boldsymbol{\gamma}$-values ranging from -2 to 2, and over 250 $\boldsymbol{\lambda}$-values ranging from 0 to 1. Per combination of $\boldsymbol{\gamma}$ and $\boldsymbol{\lambda}$ values, 10 trials, each with 100,000 random samples, were generated. For each trial, the mean and variance across the connection probabilities $\boldsymbol{p_i}$, (i.e., μ and $\boldsymbol{\sigma^2}$), the probability of each triplet motif $\boldsymbol{P}(|\boldsymbol{X}| = \boldsymbol{k})$ with $\boldsymbol{K} = \boldsymbol{6}$ (i.e., maximal number of edges in a triplet), and the respective probability expected in random network based solely on μ was calculated.

The details of the mathematical model for one trial are below:

**Algorithm 1: Mathematical model of correlated connectivity**

**Input:** simulator with degree of correlations and heterogeneity $\boldsymbol{\lambda}$ and degree of connectivity $\boldsymbol{\gamma}$.

randomly initialize shared source $\boldsymbol{S} \sim \boldsymbol{\mathcal{N}}(\boldsymbol{0}, \boldsymbol{1})$.

$\eta := 1 - \lambda$

$p_i := L(0, \gamma + \sqrt{\lambda}S, \eta)$ // $p_i$ is a vector of connection probabilities

$\mu := E(p_i)$ // $E$ denotes the expected value

$\sigma^2 := Var(p_i)$ // $Var$ denotes the variance

$K := 6$ // maximal number of edges in a triplet

**for** *k* = 0 to *K* do

  $P(k) := \binom{K}{k} E({p_i}^k (1-p_i)^{K-k})$ // probability of triplet motif with $k$ edges

  $P_{random}(k) := \binom{K}{k} \mu^k (1-\mu)^{K-k}$ // probability of triplet motif with $k$ edges in // random network

**return** $P$, $P_{random}$, $\mu$, $\sigma^2$

The deviation of motif occurrences from a random network of each triplet motif was the ratio between the means of the motif probabilities across all trials. The deviations were mapped on a grid spanned by 20 μ-values (i.e., sparsity) and 20 $\boldsymbol{\lambda}$-values (i.e., correlations & heterogeneity) and visualized by a log-space color map. Each of the 220 cell type-specific triplet combinations was mapped into the grid space. Specifically, we inferred for each combination its respective $\boldsymbol{\lambda}$-value based on the variance and mean of the connection probabilities of each combination and a lookup table of μ, $\boldsymbol{\sigma^2}$, and $\boldsymbol{\lambda}$-values as determined by the numerical simulation.

### Supplementary figures

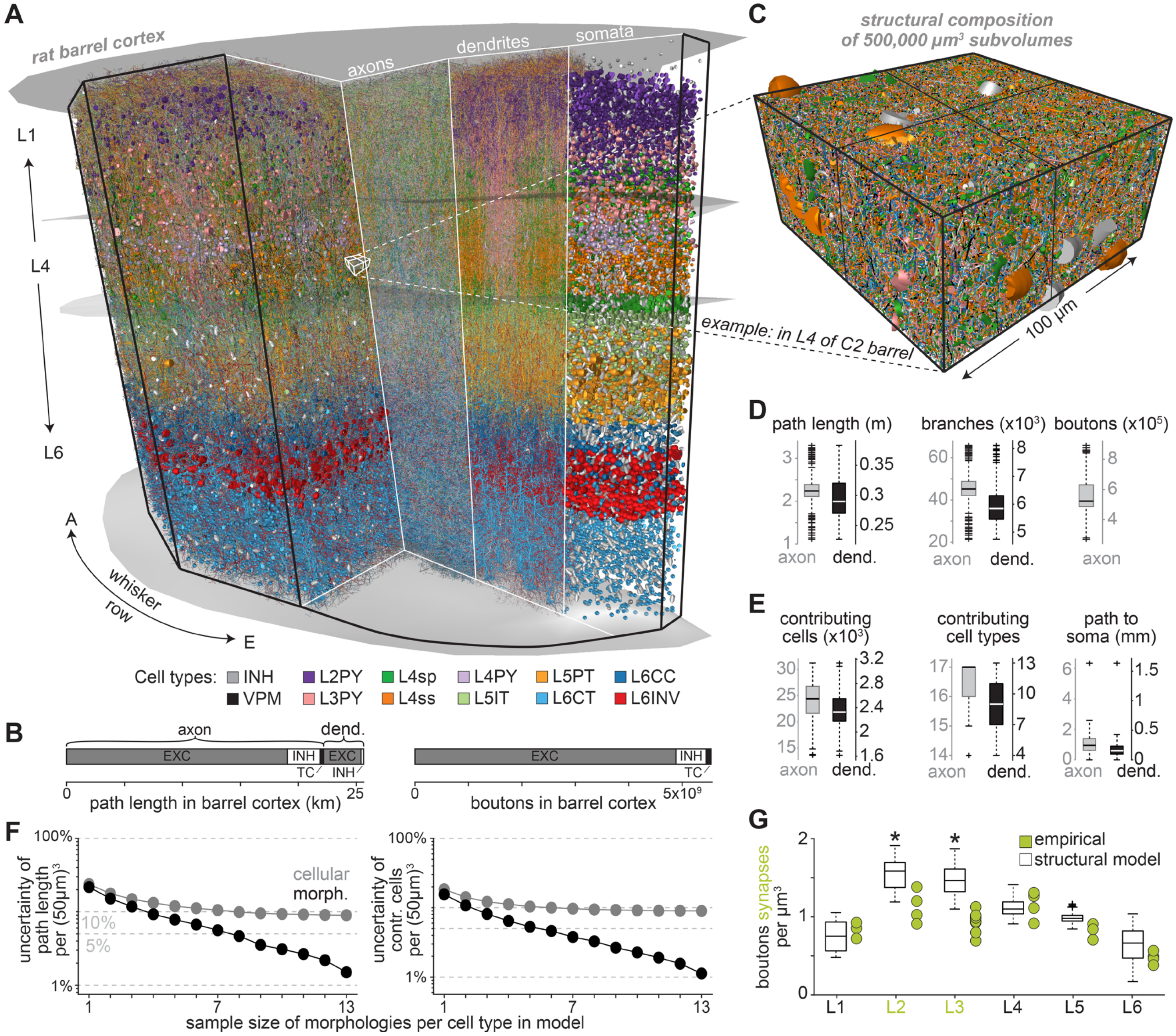


**Figure S1. Validation of the structural model of rat barrel cortex. (A)** Cross-section through the structural model of rat barrel cortex, illustrating the dense distributions of somata, dendrites and axons (fractions shown), colored by their respective cell types. **(B)** Path lengths of neuronal processes and the number of synaptic structures along them (e.g. boutons) for all excitatory (EXC) and inhibitory (INH) cortical neurons in the structural model, and thalamocortical (TC) axons from the ventral posterior medial nucleus (VPM). **(C)** Zoom-in shows one example subvolume, whose dimensions and location resemble that of the largest densely reconstructed dataset reported so far (*13*). **(D)** Path lengths, the corresponding numbers of branches, and the synaptic structures that they represent, for axons (grey) and dendrites (black) within 500,000 µm$^3$ large subvolumes (n=128) across the structural model. **(E)** Branches that are unconnected to a soma within the same subvolume (>97%) originate from >25,000 neurons (left), which reflect a diverse range of cell types (center), and whose somata are on average ~1 mm away from the subvolume (right). The 17 cell types reflect VPM, INH (subdivided by layers 1-6) and EXC neurons (subdivided as in panel A). For path to soma only maximum outlier shown. **(F)** Quantification how robustly our reverse engineering approach can predict the structural composition of each subvolume within the structural model of the rat barrel cortex. Left: robustness of the packing density of neuronal processes for each subvolume depending on the number of morphologies per cell type that was used to generate the structural model. Black markers represent the coefficients of variation (CVs) in path length per subvolume across >30,000 structural models (i.e., median across 125,000 $\mu m^3$ large subvolumes), where all structural models are based on the same average soma distribution (*42*). Grey markers represent the median CVs for the same subvolumes across structural models that were based on different empirically measured soma distributions. Right: same robustness analysis for the numbers of neurons from which these processes originate. **(G)** Predicted bouton densities per layer versus synapse density measurements in juvenile rats (*83*). The differences in layers 2 and 3 between the structural model and the empirical data (denoted by asterisks) likely reflect age differences (i.e., the data for the structural model was acquired after postnatal day P28, the empirical data at P14). Synapse densities increase particularly in the upper layers after P14 (*84*).

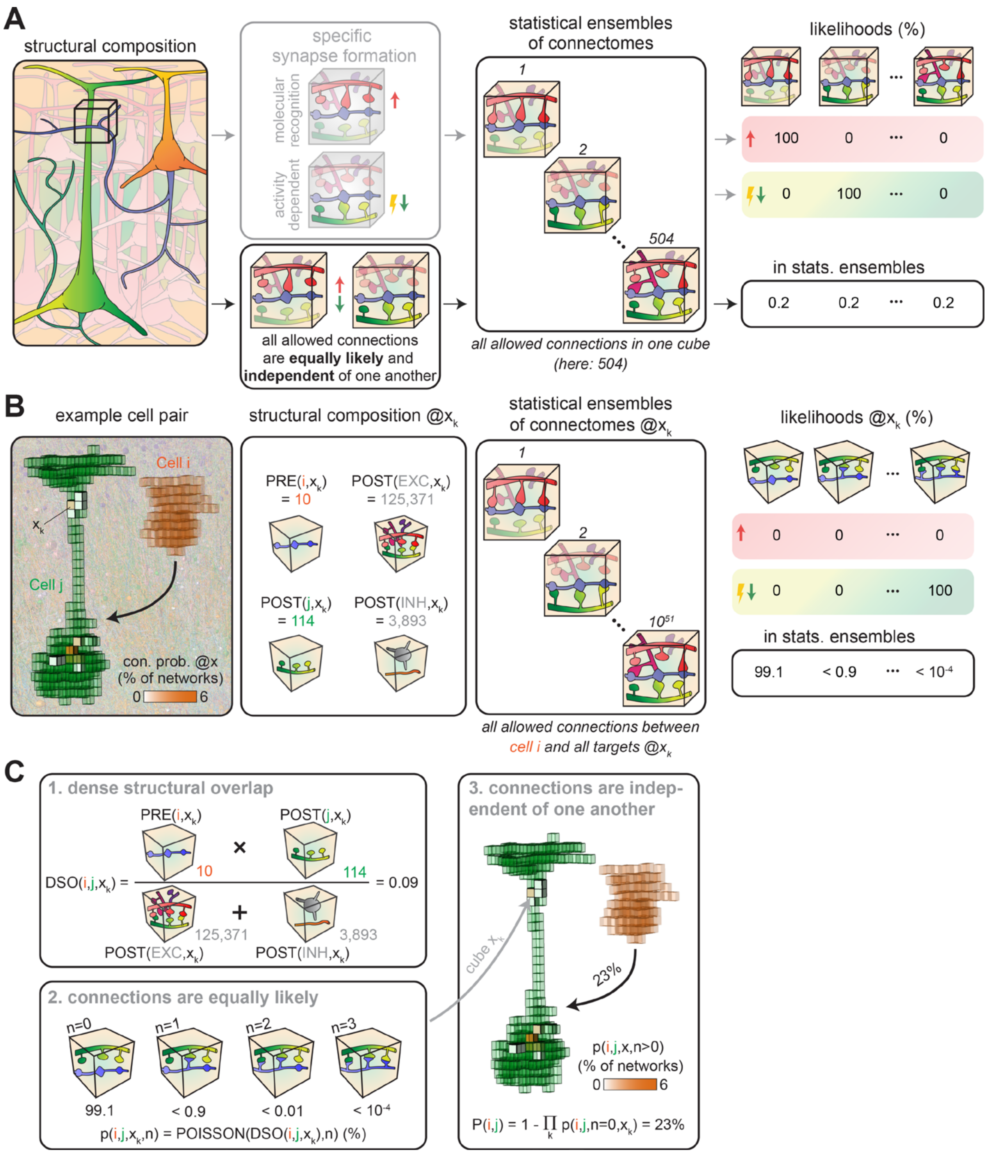


**Figure S2. Concept for testing the impact of neuron morphology on cortical network architecture. (A)** Schematically illustrated structural model of the neuropil, analogous to **Fig. 1A/B**. When each of the three axonal boutons (blue) is assigned to one of the nine spines that are present within the same example subvolume, and which represent four different neurons (green, red, pink, purple), 504 network configurations can occur that differ in how the five neurons from which these branches originate are connected to one another. If synaptic connections were formed by mechanisms that introduce wiring specificity, e.g. where axonal boutons and dendritic spines are interconnected based on their neurons' cellular or cell type identity (i.e., via molecular recognition) or electrical activity, then only one of the 504 network configurations may occur, i.e. the configuration where all boutons are connected to the spines of the same branch has a likelihood of 100%. Here, we calculate the likelihoods at which any of these network configurations could occur in the absence of synapse formation mechanisms that introduce wiring specificity. For this purpose, we assume that all allowed connections form equally likely and independently of one another. Thus, each of the 504 configurations occurs with a likelihood larger than zero. The hence generated probability distribution of neuronal network configurations (i.e., the statistical ensemble of connectomes analyzed throughout this study) reveals how neuron location and morphology could in principle impact network architecture. **(B)** Analogous to panel A, but for the neuron pair in the structural model from **Fig. 1F** and **Fig. 2A**. In one example subvolume $x_k$ at a resolution of 50 µm, ten presynaptic structures ($PRE(i, x_k)$; i.e., boutons) of $Cell\ i$ (L2PY) overlap with a total of 125,371 spines of which 114 originate from $Cell\ j$ (L5PT). In addition, the presynaptic structures overlap with 3,893 postsynaptic structures on the surfaces of inhibitory somata and dendrites (*19*). Hence, just within subvolume $x_k$ there are ~$10^{51}$ configurations by which the ten boutons of $Cell\ i$ could be connected to the available postsynaptic target structures. Thus, in 99.1% of the networks $Cell\ i$ will remain unconnected to $Cell\ j$ in subvolume $x_k$. However, the likelihood that these two neurons are connected within this subvolume by one or more synapses is larger than zero (e.g. ~$10^{-4}$% for three connections). **(C)** As reported previously (*19*), we parameterize the distributions of pre- and postsynaptic structures by calculating the quantity *dense structural overlap* ($DSO$) for any subvolume of the structural model, here illustrated between $Cell\ i$ and $Cell\ j$ in subvolume $x_k$ at a resolution of 50 μm. This quantity represents the pre- and postsynaptic structures of two neurons in an overlap volume with respect to all postsynaptic structures (see Equation 1 in **Materials and Methods**). Excitatory connections are assumed to form between boutons and spines of excitatory neurons, as well as with target sites that are distributed on the somata and dendrites of inhibitory neurons proportional to their surface areas. For inhibitory connections, target sites on excitatory and inhibitory somata and dendrites proportional to their surface areas are considered, while spines are excluded. Hence, the probability $p$ that two neurons $i$ and $j$ form $n$ synapses in a subvolume $x_k$ is given by a Poisson distribution with the quantity $DSO$ and $n$ as parameters (see Equation 2 in **Materials and Methods**). Thus, $p(i, j, x_k, n)$ denotes the percentage of networks in the statistical ensemble of connectomes in which $Cell\ i$ forms $n$ synapses with $Cell\ j$ in subvolume $x_k$. Because all connections are assumed to form independently of one another, the probability $P$ that the two neurons are connected in any of the networks by at least one synaptic connection is the product of $p$ across all overlap volumes (see Equation 3 in **Materials and Methods**).

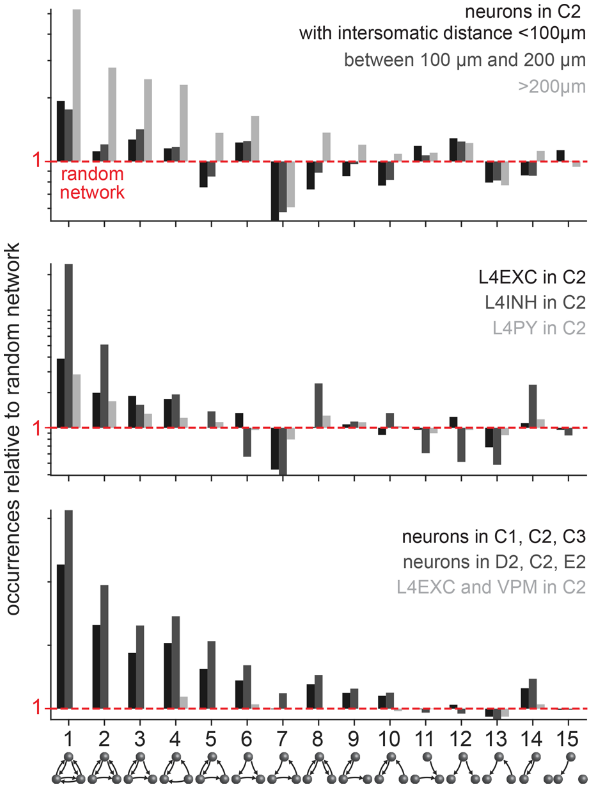


**Figure S3. Occurrences of triplet motifs depend on the grouping of neurons in the structural model of rat barrel cortex.** Ratios between motif occurrences in networks from the statistical ensemble of connectomes and random networks with the same pairwise statistics for neurons within the C2 barrel column grouped by different inter-somatic distances (top), grouped by different cell types (center), and grouped by their soma locations in different columns or VPM thalamus (bottom). Y-axis in log scale.

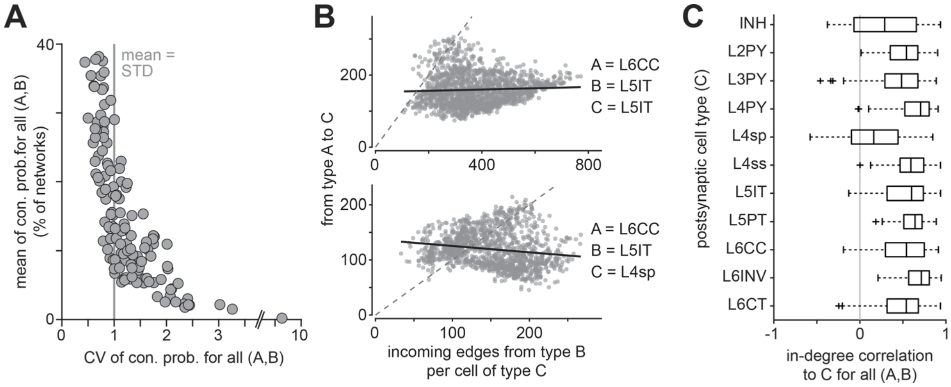


**Figure S4. Pairwise connectivity depends on the grouping of neurons in the structural model of rat barrel cortex. (A)** Means and CVs of connection probability distributions for cell type-specific groupings (10 excitatory cell types, VPM (only presynaptic), and inhibitory neurons; n=132). **(B)** Correlations between in-degree distributions for three example groupings. **(C)** In-degree correlation coefficients for different groupings (n=66 per postsynaptic cell type). All data for neurons closest to the C2 barrel column.

### Supplementary tables

| Ref. | Grouping | | | | Empirical | | | | | Predicted | | Comparison | |
|---|---|---|---|---|---|---|---|---|---|---|---|---|---|
| | Presynaptic A | | Postsynaptic B | | species | age | *in vivo* | n | P(A,B) (%) | P(A,B) MEAN (%) | P(A,B) STD (%) | dev | prctl. |
| | layer | type | layer | type | | | | | | | | | |
| (*27*) | VPM | | L4 | n/s | rat | 24-35 | yes | 40 | **43** | **42** | 29 | 0.03 | 54 |
| (*62*) | | | | | rat | adult | yes | 62 | **37** | **42** | 29 | -0.15 | 48 |
| (*27*)* | | | | | rat | 24-35 | yes | 14 | **43** | **37** | 29 | 0.20 | 60 |
| (*62*)* | | | | | rat | adult | yes | 21 | **38** | **37** | 29 | 0.03 | 55 |
| (*27*) | | | | L4sp | rat | 24-35 | yes | 24 | **42** | **43** | 27 | -0.06 | 48 |
| (*27*) | | | | L4ss | rat | 24-35 | yes | 11 | **64** | **39** | 27 | 0.90 | 78 |
| (*63*) | | | L5 | L5IT | rat | adult | yes | 18 | **17** | **27** | 22 | -0.49 | 40 |
| (*63*) | | | | L5PT | rat | adult | yes | 9 | **44** | **38** | 25 | 0.27 | 60 |
| (*63*) | | | L6 | n/s | rat | adult | yes | 11 | **9** | **17** | 18 | -0.41 | 48 |
| (*75*) | L2 | n/s | L2 | n/s | mouse | 18-21 | no | 950 | **9** | **28** | 24 | -0.79 | 29 |
| (*72*) | | | | | mouse | 18-30 | yes | 878 | **7** | **34** | 27 | -1.01 | 21 |
| (*75*) | | | L3 | n/s | mouse | 18-21 | no | 183 | **5** | **16** | 19 | -0.57 | 41 |
| (*75*) | | | L4 | n/s | mouse | 18-21 | no | 208 | **1** | **6** | 10 | -0.54 | 45 |
| (*75*) | | | L5A | n/s | mouse | 18-21 | no | 211 | **9** | **11** | 13 | -0.11 | 59 |
| (*75*) | | | L5B | n/s | mouse | 18-21 | no | 108 | **8** | **9** | 12 | -0.02 | 66 |
| (*75*) | | | L6 | n/s | mouse | 18-21 | no | 50 | **0** | **2** | 4 | -0.47 | 63 |
| (*58*) | L2/3 | n/s | L2/3 | n/s | mouse | 17-22 | no | 95 | **17** | **19** | 20 | -0.12 | 57 |
| (*66*) | | | | | rat | 17-23 | no | - | **10** | **19** | 20 | -0.46 | 44 |
| (*70*) | | | | | rat | 14-16 | no | 542 | **5** | **19** | 20 | -0.71 | 33 |
| (*71*) | | | | | mouse | 60+ | no | 112 | **2** | **19** | 20 | -0.87 | 24 |
| (*80*) | | | | | rat | adult | no | 247 | **26** | **19** | 20 | 0.35 | 70 |
| (*29*) | | | | | rat | 21-26 | no | 112 | **20** | **19** | 20 | 0.02 | 62 |
| (*69*) | | | | | mouse | adult | no | 235 | **19** | **19** | 20 | 0.00 | 60 |
| (*73*) | | | | | mouse | 21-30 | yes | 774 | **7** | **24** | 24 | -0.72 | 34 |
| (*71*) | | | L5 | L5IT | mouse | 60+ | no | 98 | **4** | **9** | 11 | -0.47 | 44 |
| (*71*) | | | | L5PT | mouse | 60+ | no | 51 | **4** | **12** | 14 | -0.61 | 36 |
| (*75*) | L3 | n/s | L2 | n/s | mouse | 18-21 | no | 182 | **12** | **13** | 15 | -0.08 | 59 |
| (*75*) | | | L3 | n/s | mouse | 18-21 | no | 513 | **19** | **20** | 19 | -0.07 | 57 |
| (*75*) | | | L4 | n/s | mouse | 18-21 | no | 170 | **2** | **10** | 12 | -0.65 | 34 |
| (*80*) | | | L5 | n/s | rat | adult | no | 29 | **55** | **8** | 10 | 4.63 | 100 |

| | | | | | | | | | | | | | |
|---|---|---|---|---|---|---|---|---|---|---|---|---|---|
| (*75*) | | | L5A | n/s | mouse | 18-21 | no | 87 | **6** | **10** | 11 | -0.36 | 48 |
| (*75*) | | | L5B | n/s | mouse | 18-21 | no | 164 | **12** | **7** | 10 | 0.56 | 80 |
| (*75*) | | | L6 | n/s | mouse | 18-21 | no | 61 | **0** | **1** | 3 | -0.34 | 73 |
| (*75*) | L4 | n/s | L2 | n/s | mouse | 18-21 | no | 208 | **12** | **14** | 16 | -0.11 | 59 |
| (*67*) | | | L2/3 | n/s | rat | 17-23 | no | 64 | **15** | **18** | 18 | -0.15 | 55 |
| (*29*) | | | | | rat | 21-26 | no | 50 | **20** | **18** | 18 | 0.12 | 63 |
| (*75*) | | | L3 | n/s | mouse | 18-21 | no | 172 | **15** | **21** | 19 | -0.34 | 46 |
| (*80*) | | | | | rat | adult | no | 25 | **28** | **21** | 19 | 0.35 | 69 |
| (*61*) | | | L4 | n/s | rat | 14-21 | no | 89 | **6** | **20** | 19 | -0.73 | 31 |
| (*75*) | | | | | mouse | 18-21 | no | 1046 | **24** | **20** | 19 | 0.21 | 65 |
| (*68*) | | | L5A | n/s | rat | 17-23 | no | - | **14** | **12** | 13 | 0.19 | 70 |
| (*75*) | | | | | mouse | 18-21 | no | 276 | **12** | **12** | 13 | 0.01 | 65 |
| (*75*) | | | L5B | n/s | mouse | 18-21 | no | 136 | **8** | **6** | 7 | 0.30 | 70 |
| (*75*) | | | L6 | n/s | mouse | 18-21 | no | 93 | **3** | **2** | 4 | 0.29 | 79 |
| (*59*) | | L4PY | L4 | L4PY | rat | adult | no | 528 | **4** | **15** | 15 | -0.70 | 34 |
| (*28*) | | L4sp | | L4sp | rat | 21-35 | no | 24 | **21** | **23** | 15 | -0.17 | 48 |
| (*65*) | | L4ss | | L4ss | rat | 12-15 | no | 94 | **26** | **26** | 22 | -0.04 | 55 |
| (*78*) | | | | | rat | 13-15 | no | 146 | **36** | **26** | 22 | 0.42 | 68 |
| (*28*) | | | | | rat | 21-35 | no | 24 | **21** | **26** | 22 | -0.25 | 48 |
| (*80*) | L5 | n/s | L3 | n/s | rat | adult | no | 29 | **3** | **6** | 11 | -0.23 | 64 |
| (*71*) | | | L5 | n/s | mouse | 60+ | no | 150 | **0** | **10** | 11 | -0.89 | 13 |
| (*76*) | | | | | rat | 14-16 | no | 500 | **10** | **10** | 11 | -0.01 | 62 |
| (*79*) | | | | | rat | 14-16 | no | 1450 | **12** | **10** | 11 | 0.16 | 68 |
| (*80*) | | | | | rat | adult | no | 163 | **9** | **10** | 11 | -0.08 | 59 |
| (*71*) | | L5IT | L2/3 | n/s | mouse | 60+ | no | 98 | **0** | **9** | 13 | -0.67 | 23 |
| (*11*) | | | L5 | L5IT | mouse | 14-17 | no | 118 | **5** | **12** | 10 | -0.66 | 32 |
| (*71*) | | | | | mouse | 60+ | no | 66 | **0** | **12** | 10 | -1.17 | 5 |
| (*11*) | | | | L5PT | mouse | 14-17 | no | 86 | **19** | **15** | 13 | 0.28 | 69 |
| (*71*) | | | | | mouse | 60+ | no | 36 | **0** | **15** | 13 | -1.21 | 4 |
| (*71*) | | L5PT | L2/3 | n/s | mouse | 60+ | no | 51 | **0** | **2** | 5 | -0.47 | 53 |
| (*11*) | | | L5 | L5IT | mouse | 14-17 | no | 86 | **5** | **8** | 9 | -0.41 | 47 |
| (*71*) | | | | | mouse | 60+ | no | 36 | **0** | **8** | 9 | -0.96 | 11 |
| (*11*) | | | | L5PT | mouse | 14-17 | no | 225 | **7** | **17** | 13 | -0.73 | 29 |
| (*71*) | | | | | mouse | 60+ | no | 12 | **0** | **17** | 13 | -1.25 | 5 |
| (*30*) | | | | | rat | 14-16 | no | 3446 | **13** | **17** | 13 | -0.32 | 46 |
| (*16*) | | | | | rat | 12-20 | no | 8050 | **12** | **17** | 13 | -0.39 | 43 |

| | | | | | | | | | | | | | |
|---|---|---|---|---|---|---|---|---|---|---|---|---|---|
| (*75*) | L5A | n/s | L2 | n/s | mouse | 18-21 | no | 209 | **4** | **11** | 15 | -0.45 | 50 |
| (*75*) | | | L3 | n/s | mouse | 18-21 | no | 89 | **2** | **9** | 14 | -0.50 | 48 |
| (*75*) | | | L4 | n/s | mouse | 18-21 | no | 275 | **1** | **8** | 11 | -0.67 | 32 |
| (*75*) | | | L5A | n/s | mouse | 18-21 | no | 934 | **19** | **15** | 13 | 0.32 | 69 |
| (*75*) | | | L5B | n/s | mouse | 18-21 | no | 175 | **8** | **7** | 8 | 0.15 | 68 |
| (*75*) | | | L6 | n/s | mouse | 18-21 | no | 158 | **3** | **3** | 4 | 0.08 | 69 |
| (*75*) | L5B | n/s | L2 | n/s | mouse | 18-21 | no | 104 | **1** | **3** | 7 | -0.26 | 73 |
| (*75*) | | | L3 | n/s | mouse | 18-21 | no | 167 | **2** | **3** | 7 | -0.18 | 72 |
| (*75*) | | | L4 | n/s | mouse | 18-21 | no | 137 | **1** | **2** | 5 | -0.35 | 64 |
| (*75*) | | | L5A | n/s | mouse | 18-21 | no | 174 | **2** | **6** | 8 | -0.53 | 45 |
| (*74*) | | | L5B | n/s | mouse | 14-37 | no | 269 | **9** | **13** | 13 | -0.31 | 48 |
| (*75*) | | | | | mouse | 18-21 | no | 555 | **7** | **13** | 13 | -0.47 | 41 |
| (*75*) | | | L6 | n/s | mouse | 18-21 | no | 100 | **7** | **6** | 9 | 0.09 | 69 |
| (*75*) | L6 | n/s | L2 | n/s | mouse | 18-21 | no | 50 | **0** | **0** | 1 | -0.13 | 96 |
| (*75*) | | | L3 | n/s | mouse | 18-21 | no | 64 | **0** | **0** | 3 | -0.18 | 91 |
| (*75*) | | | L4 | n/s | mouse | 18-21 | no | 94 | **0** | **1** | 4 | -0.31 | 75 |
| (*75*) | | | L5A | n/s | mouse | 18-21 | no | 160 | **1** | **3** | 6 | -0.48 | 58 |
| (*75*) | | | L5B | n/s | mouse | 18-21 | no | 100 | **2** | **6** | 8 | -0.48 | 49 |
| (*60*) | | | L6 | n/s | rat | 14-21 | no | 102 | **2** | **11** | 13 | -0.71 | 34 |
| (*75*) | | | | | mouse | 18-21 | no | 532 | **3** | **11** | 13 | -0.64 | 38 |
| (*77*) | | | | | rat | adult | no | 27 | **4** | **11** | 13 | -0.57 | 42 |
| (*64*) | L6A | L6CC | L6A | L6CT | mouse | 20-33 | no | 43 | **9** | **14** | 10 | -0.51 | 36 |
| (*64*) | L6A | L6CT | L6A | L6CC | mouse | 20-33 | no | 40 | **0** | **12** | 12 | -1.00 | 12 |

**Table S1. Predicted versus empirical connectivity data.** An asterisk (*) after the reference denotes connectivity measurements within the septum; age is given in postnatal days; n denotes the number of pairs tested; P(A,B) denotes the connection probability of type A to type B, either measured or predicted by the statistical ensemble of connectomes (% of networks) as the mean or standard deviation (STD). Deviation between empirically observed and predicted connection probability: dev = (P(A,B) empirical – mean of P(A,B) predicted) / (STD of P(A,B) predicted). prctl denotes percentage of predicted connection probabilities that are lower or equal than P(A,B) empirical (e.g., prctl = 0: none of the predicted connection probabilities are lower or equal than P(A,B) empirical; prctl = 100: all predicted connection probabilities are lower or equal than P(A,B) empirical). Correlation between predictions and empirical data: R=0.71 ($p<10^{-5}$, n=32 for rat); R=0.55 ($p=10^{-5}$, n=57 for mouse)

| Inter-somatic distance (µm) | Connection probability L5PT to L5PT (%) | | | |
|---|---|---|---|---|
| | empirical | predicted | | |
| | | median | 25th percentile | 75th percentile |
| 18 | 22 | 23 | 15 | 34 |
| 53 | 17 | 21 | 13 | 31 |
| 88 | 15 | 18 | 10 | 28 |
| 123 | 13 | 15 | 7 | 24 |
| 158 | 10 | 11 | 4 | 19 |
| 193 | 10 | 7 | 2 | 14 |
| 228 | 7 | 4 | 1 | 10 |
| 263 | 4 | 2 | 0 | 7 |
| 298 | 7 | 1 | 0 | 6 |

**Table S2. Predicted versus empirical connectivity data.** Inter-somatic distance-dependent connection probabilities between L5PTs. Empirical data from (*30*).

| Inter-somatic distance (µm) | Connection probability L2/3EXC to L2/3EXC (%) | | | |
|---|---|---|---|---|
| | empirical | predicted | | |
| | | median | 25th percentile | 75th percentile |
| 20 | 21 | 30 | 15 | 48 |
| 60 | 17 | 25 | 11 | 42 |
| 100 | 13 | 17 | 7 | 32 |
| 140 | 14 | 11 | 3 | 24 |

**Table S3. Predicted versus empirical connectivity data.** Inter-somatic distance-dependent connection probabilities between EXC neurons in layers 2/3. Empirical data from (*58*).
